# Migration and division in cell monolayers on substrates with topological defects

**DOI:** 10.1101/2022.12.22.521493

**Authors:** Kurmanbek Kaiyrbekov, Kirsten Endresen, Kyle Sullivan, Zhaofei Zheng, Yun Chen, Francesca Serra, Brian A. Camley

## Abstract

Collective movement and organization of cell monolayers are important for wound healing and tissue development. Recent experiments highlighted the importance of liquid crystal order within these layers, suggesting that +1 topological defects have a role in organizing tissue morphogenesis. We study fibroblast organization, motion and proliferation on a substrate with micron-sized ridges that induce +1 and −1 topological defects using simulation and experiment. We model cells as self-propelled deformable ellipses that interact via a Gay-Berne potential. Unlike earlier work on other cell types, we see that density variation near defects is not explained by collective migration. We propose instead that fibroblasts have different division rates depending on their area and aspect ratio. This model captures key features of our previous experiments: the alignment quality worsens at high cell density and, at the center of the +1 defects, cells can adopt either highly anisotropic or primarily isotropic morphologies. Experiments performed with different ridge heights confirm a new prediction of this model: suppressing migration across ridges promotes *higher* cell density at the +1 defect. Our work enables new mechanisms for tissue patterning using topological defects.

## I. INTRODUCTION

Monolayers of cells in multicellular organisms cooperate to transmit forces in embryogenesis, act as a barrier, and perform many more essential functions [1]. These cells often have long axes locally aligned with each other – i.e. they have local nematic order akin to liquid crystals [2–5]. Deviations from perfect nematic alignment can occur as topological defects. In 2D, defects are points where following cell orientation for a complete cycle around the defect leads to a rotation in orientation Δ*θ* = 2*πq*, where the topological charge *q* is integer (*q* = ±1 shown in Fig. 1a) or half integer. Topological defects are biologically relevant: they can drive cell death and extrusion [6], cell dynamics [7], tissue branching in regeneration [8], and growth [9]. These defects can also reorganize cell density. Recent experiments with monolayers of various cell types show cells tend to congregate at positive defects and disperse at negative ones [9–11], though this is not universal to all cell types [6, 12]. Congregation at positive defects can result in increased density [10], creation of new layers of cells[11], or growth of mounds of cells [9]. In all these examples, accumulation at defects with positive topological charge and depletion at defects with negative topological charge is driven by collective migration of cells. Here, we want to understand how we can control cell density, shape, and cell orientation by exploiting the topological properties of cell monolayers. We use our earlier-developed system of NIH 3T6 fibroblasts on a substrate with micron-scale ridges [12]; fibroblasts align their long axes along the ridges. When we impose a +1 or −1 defect pattern on the ridge (Fig. 1a), this forces cells to take up this defect pattern (Fig. 1b). We also see enhanced density at +1 defects and decreased density at −1 defects. However, we find that system lacks large scale collective migration that could drive density variations at defects. We use simulations and experiments to show that shape-dependent division is sufficient to cause density variations at defects. This is a qualitatively new way to pattern cells using topological defects.

**FIG. 1.**
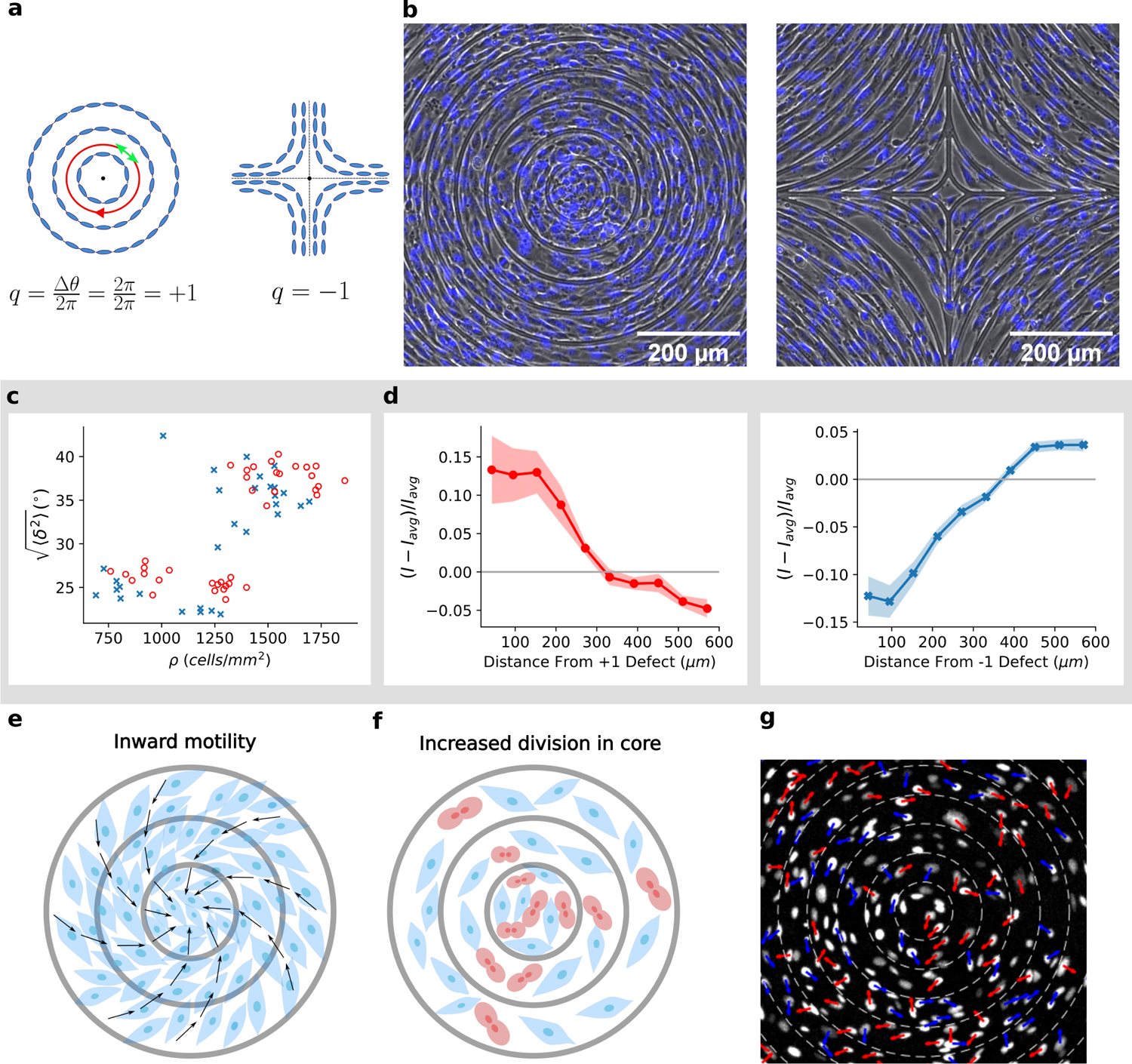
Experimental results, data in gray box replotted from [12]. **a**, Topological defect of charge +1 (left), the nematic director shown by a green arrow rotates by 2π as it circles around defect (shown by red path). Similarly −1 defect (right). **b**, Phase contrast image of 3T6 cells in the vicinity of a positive defect (left) and negative defect (right), overlaid with fluorescent image of nuclei stained with Hoechst 33342. The spacing between ridges is 60 *μm* and ridges are 1.5 *μm* tall. **c** Root mean square deviation (RMSD) 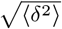 from ideal alignment for positive defects (red circles, n=39 defects) and negative defects (blue crosses, n=30 defects). Each data point corresponds to observation of cells near one defect, averaged over the cells **d**, Density of fibroblasts as a function of distance from center of positive (left) and negative (right) defects. Density is determined from nuclear fluorescence (NucRed Live 647 or Hoechst 33342; *Methods*). Shown is the deviation from the sample’s average intensity *I*_avg_, normalized by *I*_avg_. Curve is averaged over many different patterns, with final densities ranging over 600-2000 cells/mm^2^ for +1 defects (*n* = 20) and –1 defects. (*n* = 28). Colored regions indicate ±1 standard error. (**e, f**): Possible modes of density increase: 1. Net inward movement of cells (**e**); black arrows represent the direction of movement of cells 2. Cell division rate differences (**f**) where there are relatively more cells dividing (shown in red) close to core of the defect. **g**, Experimental measurement of fibroblast velocity. Fibroblast displacement direction over 1 h shown by arrows. The arrows are colored blue if the component of the net displacement parallel to the ridges is in the counter-clockwise direction and red if clockwise. Tracks are shown in Supplementary Movie 1.

## II. RESULTS

### A. Experiments: cell density increases at +1 defects, but likely not through migration

We use photolithography to create a substrate with 1.5-*μ*m-high ridges in a pattern chosen to induce +1 and −1 topological defects (Fig. 1a-b). We seed 3T6 fibroblast cells on the substrates and observe their behavior as they proliferate. Our earlier work [12] discovered three key features, which we reproduce in Fig. 1c-d (gray box): 1) the fibroblasts’ long axes follow the ridges, 2) the degree of deviation from the ridges increases as cells are increasingly packed past confluence, and 3) fibroblast density is increased relative to the rest of the monolayer at the center of the +1 defect pattern and relatively decreased at the center of the −1 defect pattern.

It is apparent in Fig. 1b that the long axis of the fibroblasts follows the direction imposed by the ridges – though imperfectly. We measure the quality of cell alignment via the root mean square deviation of cell long axis orientation *θ* from expected alignment *θ_e_* (see Methods), 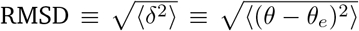. Cells at subconfluent density *ρ* < 1000 cells/mm^2^ are relatively well-aligned, but the quality of alignment decreases at larger densities (Fig. 1c).

We also measure cell density via nuclear fluorescence *I*(*r*), as a function of the distance *r* from the pattern center. Fibroblasts have an elevated density close to the +1 defect and low density at the −1 defect (Fig. 1d).

How does the topographic pattern change density at the defect core? Previous studies on myoblasts, neural progenitor cells, and myxobacteria argued density differences at defects arise due to collective motion of cells [9–11], including dramatic inspiraling migration [9]. We sketch this broad mechanism in Fig. 1e. However, increased density at the +1 defect could also arise from higher proliferation rate of cells near the +1 defect (Fig. 1f). If migration were driving the increase in density in our experiments as in other cell types, we would expect significant inward migration towards +1 defects and away from −1 defects. We find instead that fibroblasts primarily move azimuthally around the +1 defect, with short-range correlation of velocities, but no broad inward flow (Fig. 1g and Supplementary Movie 1). Given the lack of collective directed motion, we hypothesize cell division rate differences are the driving factor of our observed density differences, and we develop a model with this assumption.

### B. Simulations reproduce experimental alignment and movement patterns

We model spindle-shaped fibroblasts as self-propelled elliptical particles. Cell *i* has semi-major axis length *a_i_* and semi-minor axis length *b_i_* (Fig. 2a). Our model includes cell motility, cell-cell interactions, and cell division: we give a full description in the Methods, and a brief summary here. At every step in the model, cells are chosen randomly to either move, rotate or alter one of their axis lengths (Fig. 2b) and we accept this move with a probability depending on the change in system energy. The energy is determined by cell shape, cell-cell interactions, cell-substrate interactions, and cell polarity. One attempt for each cell is a “Monte Carlo Step” (MCS); we calibrate parameters so 100 MCS is 1.5 minutes of experimental time.

**FIG. 2.**
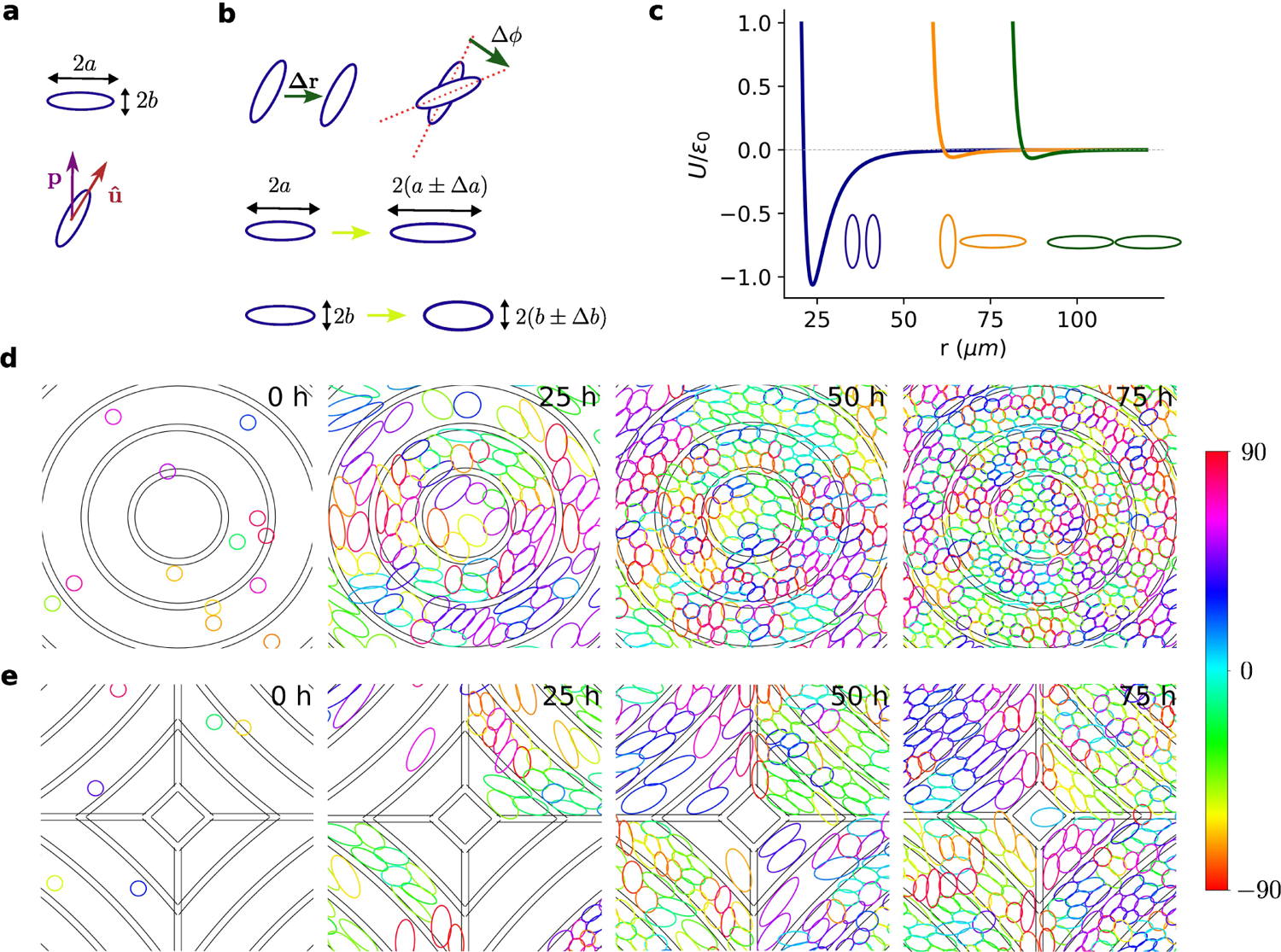
Model details and time evolution of simulation. **a**, Cells are modeled as elliptical particles with the *i*^th^ cell having semi-major axis length *a_i_* and semi-minor axis length *b_i_*. The orientation of a cell is represented by a unit vector *û* which points along the long axis of the ellipse. The polarity vector ***p*** denotes preferred direction of motion. **b** List of possible move at each Monte-Carlo step: displacement by Δ*r*, rotation by Δ*ϕ*, change of semi-major axis length ±Δ*α* or change of semi-minor axis length ±Δ*b*. **c** Cell pairs interact via a modified Gay-Berne potential. The potential is weakly attractive at long separations, strongly repulsive at shorter distances. Parallel alignment of long axes (blue) of pairs of cells is preferred over other configurations (orange, green). **d**,**e**: Time evolution of the simulation at the core of +1 **d** and −1 **e** defect from start (left) to T=75h (right). Cells are colored according to the angle they make with the x axis. Small concentric regions between two consecutive dark rings represents a ridge. These are zoomed-in views, showing a subsection of the the whole simulation box (full simulation box is 1200 *μ*m×1200 *μ*m). See also Supplementary Movies 2, 3.

The cell shape energy models cells tending to keep their area *A* = *πab* and aspect ratio *AR* = *a/b*; we set *AR*_pref_ = 4, *A*_pref_ = 1400*μm*^2^ to match experiment [4]. Deviations from preferred aspect ratios and areas have energy cost proportional to *k_AR_* and *k_Area_* respectively, setting the “stiffnes” of the cell’s shape. We also include a core energy that prevents indefinite squeezing of cells. Cells interact with one another via a modified Gay-Berne [13–16] potential widely used in liquid crystal simulations (Fig. 2c); this energy promotes cells having their long sides adjacent to one another, inducing nematic order. Cell-ridge overlap is penalized with energy cost equal to the product of ridge strength *k_r_* and fraction of the cell overlapping with the ridge. We argue that ridge strength reflects the ridge height in experiments.

Crawling eukaryotic cells are chemically and mechanically polarized [17]; we summarize this polarity by a vector ***p*** indicating the direction the cell prefers to move (Fig. 2a). Fibroblasts move along their long axis [4]. We add a motility energy that encourages motion along ***p*** and the long axis in a direction **Π** = **û**(**û** · **p**), where ***û*** is the long axis of the cell. Fibroblast polarity occasionally flips direction [4], which we model by stochastically reversing **p** with average flip period of ~ 2.5 hours. We also assume cells tend to align to their past displacement [18–20], leading to some coordination between cell velocities, as in Fig. 1g. In between polarity flips, **p** obeys a rule proposed by Szabo et. al. [21] where after *t* MCS, we update the polarity vector for each cell as ***p**_t_* = (1 – 1/*τ*_pol_)***p***_*t*–1_ + **Δ*r*** where **Δ*r*** = (Δ*x*, Δ*y*) is the proposed displacement, ***p***_*t*–1_ is polarity for that cell at time-step *t* – 1 and *τ*_pol_ is the polarity decay timescale measured in MCS. Polarity thus reorients toward the most recent displacement, promoting cells crawling persistently and coherently [19]. This is disrupted by polarity flipping.

Within our model, we seed initially small, circular cells at density of *ρ*_init_ ~ 70 cells/mm^2^ randomly in our periodic simulation box, and then choose cells to divide at a rate set by the experimental growth curve (Fig. S1). The probability that cell *i* is selected for division is *p_i_*, given by Eq. 1, discussed in more detail in the next section.

We show a typical simulation in Fig. 2d, e and Supplementary Movies 2, 3. The −1 defect is constructed by using the periodic boundary condition (Fig. S2). We track simulated cells’ RMSD from perfect alignment with the ridge pattern 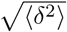 as cells proliferate, and observe that RMSD increases as cells reach higher density (Fig. 3a). This is consistent with our experiments (Fig. 1c). In our model, alignment is also controlled by ridge strength *k_r_* – larger *k_r_* decreases RMSD (Fig. 3a), though this effect saturates for *k_r_* > 100. (*k_r_* values are relative to the effective temperature *T*; *Methods*.) These results are simply a consequence of packing anisotropic deformable cells, and do not require cell motility or a particular division mechanism (Fig. S3). Sensitivity to ridges may decrease at large densities because cells become less anisotropic – hence less able to coherently align. The experimental analog to ridge strength is ridge height above the substrate. We vary ridge height experimentally (Fig. 3b), finding weak effects on RMSD, suggesting experiments are near the limit where increasing *k_r_* has diminishing returns on alignment.

**FIG. 3.**
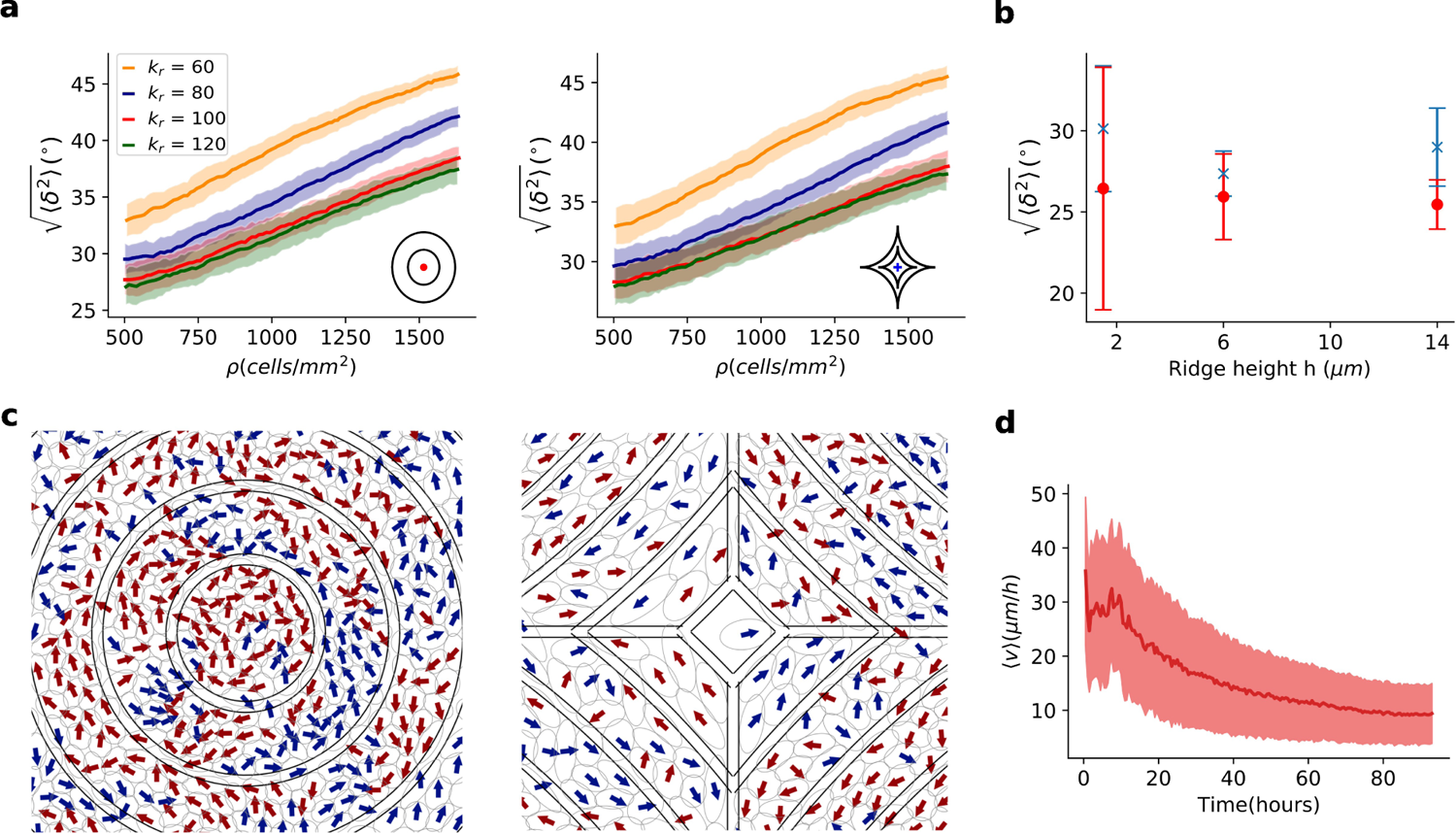
Alignment and migration patterns of cells. **a** Root Mean Square Deviation (RMSD) 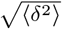 of cell alignment from an expected orientation as a function of cells density for various ridge strengths. Shaded area shows standard deviation of RMSDs of 100 simulations. **b** Ridge height dependence of RMSDs with respect to +1 defect (red) and −1 defect (blue). Error bars are standard deviations. **c** Directions of cell motion shown by arrows. The arrows are colored blue if the net displacement is in counter-clockwise direction and by red if clockwise. **d** Average cell speeds during simulation. Averaging is done over cells and shaded area is standard deviation.

Simulated cell motion resembles experimental trajectories (Fig. 1g). We see locally correlated, primarily azimuthal motion without overall coherent direction (Fig. 3c). Average cell speeds slow over time as the monolayer becomes more densely packed, broadly consistent with past measurements [4].

Our model recapitulates experimental cell motion and alignment. Can we understand the increase in density at +1 defects and decrease in density at −1 defects?

### C. Shape dependent division is sufficient to drive density variations at defects

Higher density of cells near +1 and lower density at −1 defects could arise from cell migration or cell proliferation (Fig. 1e,f). Given the lack of clear inward migration (Fig. 1g), we hypothesize cell proliferation rates are different near defects. One possible reason for this difference is that the cell shapes near the defects differ. Past work on confinement and stretching experiments with endothelial and smooth muscle cells demonstrated that uniform or uniaxial compression suppresses division rate [22, 23]. We thus propose a model where larger cells and more isotropic cells are more likely to divide. We set the probability of cell *i* to be selected to divide as

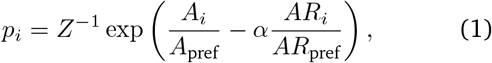

where the shape sensitivity *α* tunes how sensitive division is to cell shape and *Z* is a normalization factor. When *α* = 0, cells with the biggest area are the most likely to divide independent of AR; as *α* increases from 0 to 2 more isotropic cells (*AR* → 1) with large areas become more likely to divide (Fig. 4a). Since the cells are more likely to be isotropic within the inner rings of the +1 defect, and isotropic cells more likely to divide, shape-dependent division can potentially drive the density variations.

**FIG. 4.**
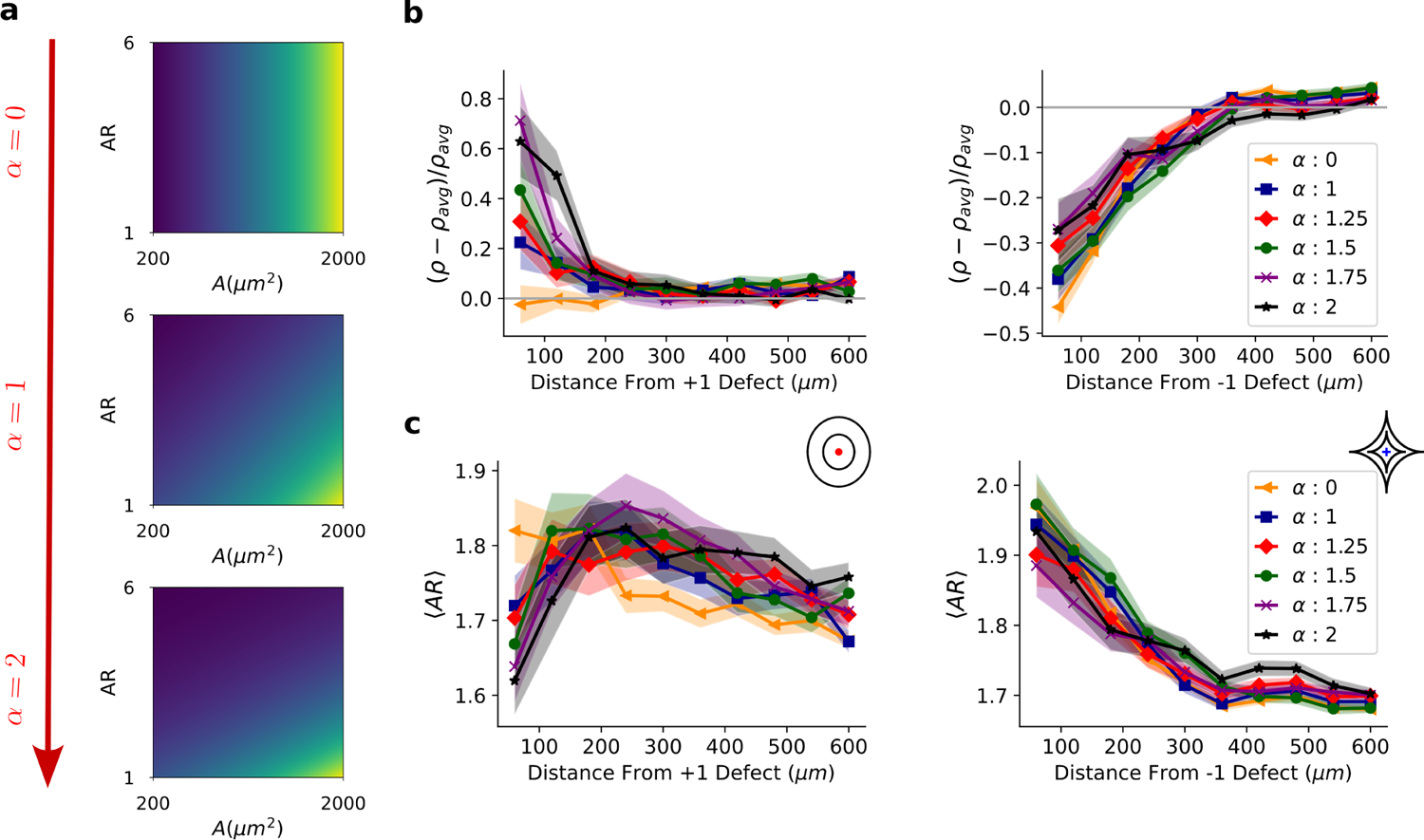
Effect of cell shape dependent division mechanism on density and aspect ratio profiles. **a**, Division probability (Eq. 1) as a function of shape sensitivity *α*, aspect ratio *AR*, and area *A*. Brighter colors (more yellow) indicate larger probability. Here for demonstration, we assume cells have uniform aspect ratios and areas. **b** Density and **c** aspect ratios as a function of distance from core of defect for various *α*. +1 defect on the left and −1 defect on the right. Colored regions represent standard errors of the mean over 100 simulations. *k_r_* = 120 in this figure.

To test whether shape-dependent division is sufficient to reproduce density variation, we vary sensitivity to shape *α* and observe density changes near defects. We show the change in density relative to the whole-system average density *ρ*_avg_ in Fig. 4b – analogous to experiments in Fig. 1d. When division probability is independent of shape (*α* = 0) density does not strongly depend on distance from the +1 defect, but density is below its average value near the −1 defect. (Similar results are found when cells are randomly selected to divide: see Fig. S4.) When we increase *α* → 2, making isotropic cells more likely to divide, relative density at the +1 defect center grows significantly. There is also a slight increase in density near the −1 defect for larger *α* (Fig. 4b), but the normalized density deviation from the average remains negative. Cell shapes also change. As *α* → 2, we see that cells near +1 defect become more isotropic than their surrounding cells. On the other hand, cells at the core of the −1 defect are always more elongated than those further away (Fig 4c). These patterns are consistent with our expectation that, when *α* = 2, isotropic cells that are more likely to divide are near +1 defects and that decreased aspect ratios allow more cells to pack near +1 defects, creating a positive feedback loop between cell shape and density.

Increased density at the +1 defect is made more prominent by cell motility, but can also be seen without it (*k*_move_ = 0; Fig. S5). In the absence of motility, cells rarely cross ridges, so the ~ 40% of simulations that start with no cell in the inner ring still have low density in the inner ring. Even though density at the +1 defect may be high in the other 60% of simulations, the large fraction of simulations with zero or low density at the core means the overall density increase at the +1 defect is weaker in the absence of motility.

### D. Simulations and experiments have high variability of cell shapes

When *α* = 2 in our simulation, cells near the defect are more isotropic than those further away (Fig. 4c), but this is highly variable between simulation runs. Increasing α from 0 to 2 increases the number of simulations with isotropic cells at the core – but there are many cases with anisotropic cells (Fig. 5a). We also see similar variability experimentally. In both experiments and simulations, we observe both patterns with elongated cells at the +1 defect center (Fig. 5b) and isotropically packed cells at the +1 defect (Fig. 5c).

**FIG. 5.**
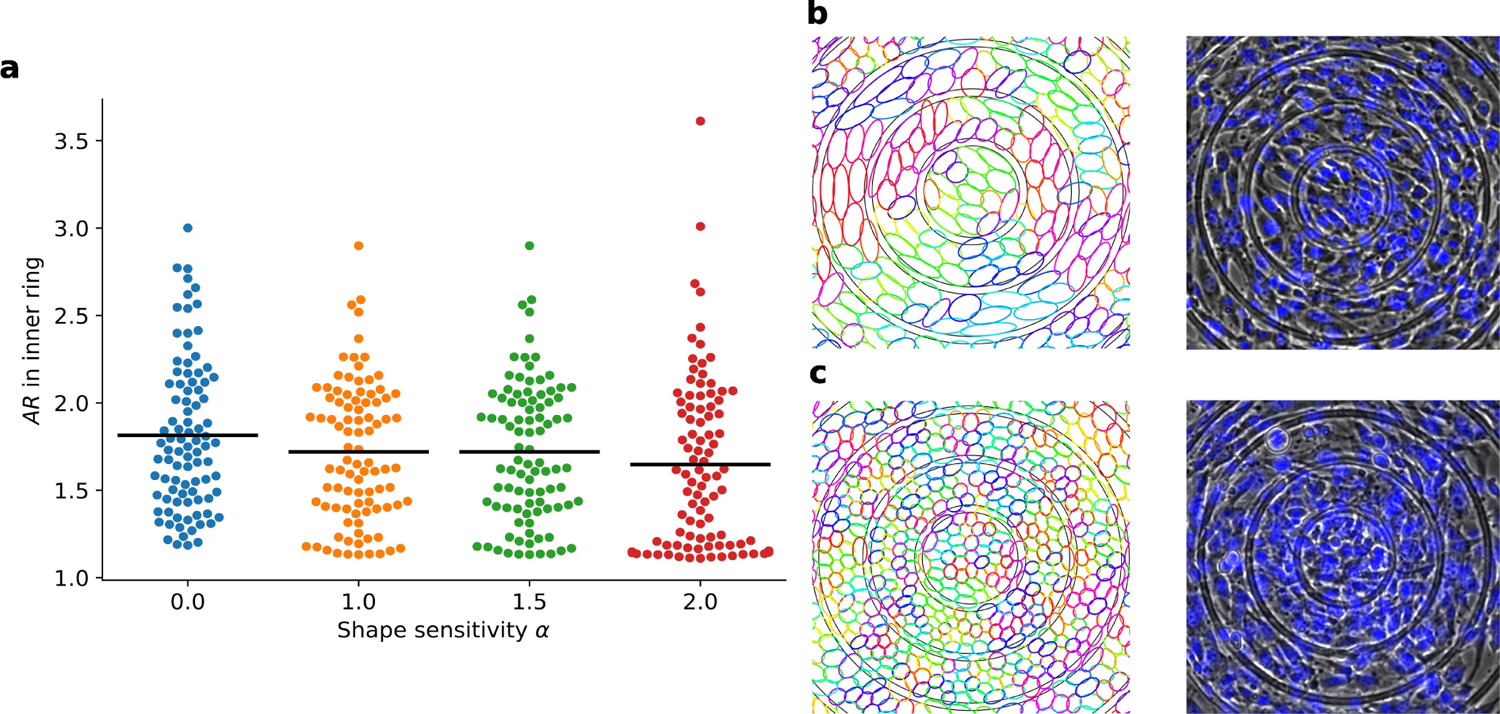
Different cell shapes at the core of +1 defect. **a**, Swarmplot of average cell aspect ratios at the innermost ring. Each dot is the average AR in the inner ring at the end of one simulation. There are 100 simulations for each sensitivity of cell division probability to shape *α*. **b**, **c** Simulation (left) and experiment (phase contrast microscopy with overlapped fluorescent nuclei, right) that end up with anisotropic (**b**) or isotropic cells (**c**) at the core.

### E. Simulations and experiments agree that increasing ridge height enhances density variations

How can we test our idea that division is driving density variations? If collective migration is the cause of density variations, we expect preventing cells from crossing ridges to suppress density changes [24]. On the other hand, if shape-dependent division drives density differences, then density variations should increase if we constrain cell movement across the ridges. At first, in our simulation, we increase ridge strength *k_r_* from 60 to 120 to reduce ridge crossing (illustrated in Supplementary Movies 4,5), finding that the number of cells overlapping with ridges drops dramatically as ridge strength grows (Fig. 6a, b). The decrease in ridge crossing is accompanied with marked changes in density (Fig. 6c) and aspect ratio (Fig. S6) near defects. We see a relatively uniform density near the +1 defect for weak ridge strength *k_r_* = 60, but we see much-increased relative density at the +1 defect as *k_r_* → 120 (Fig. 6c). We see the opposite trend near −1 defects, with relative density decreasing with *k_r_*, though this saturates as *k_r_* ≈ 80 - 120. Thus, simulations predict that preventing cell crossings enhances density at the core of +1 defect – as expected if cell division drives the density increase.

**FIG. 6.**
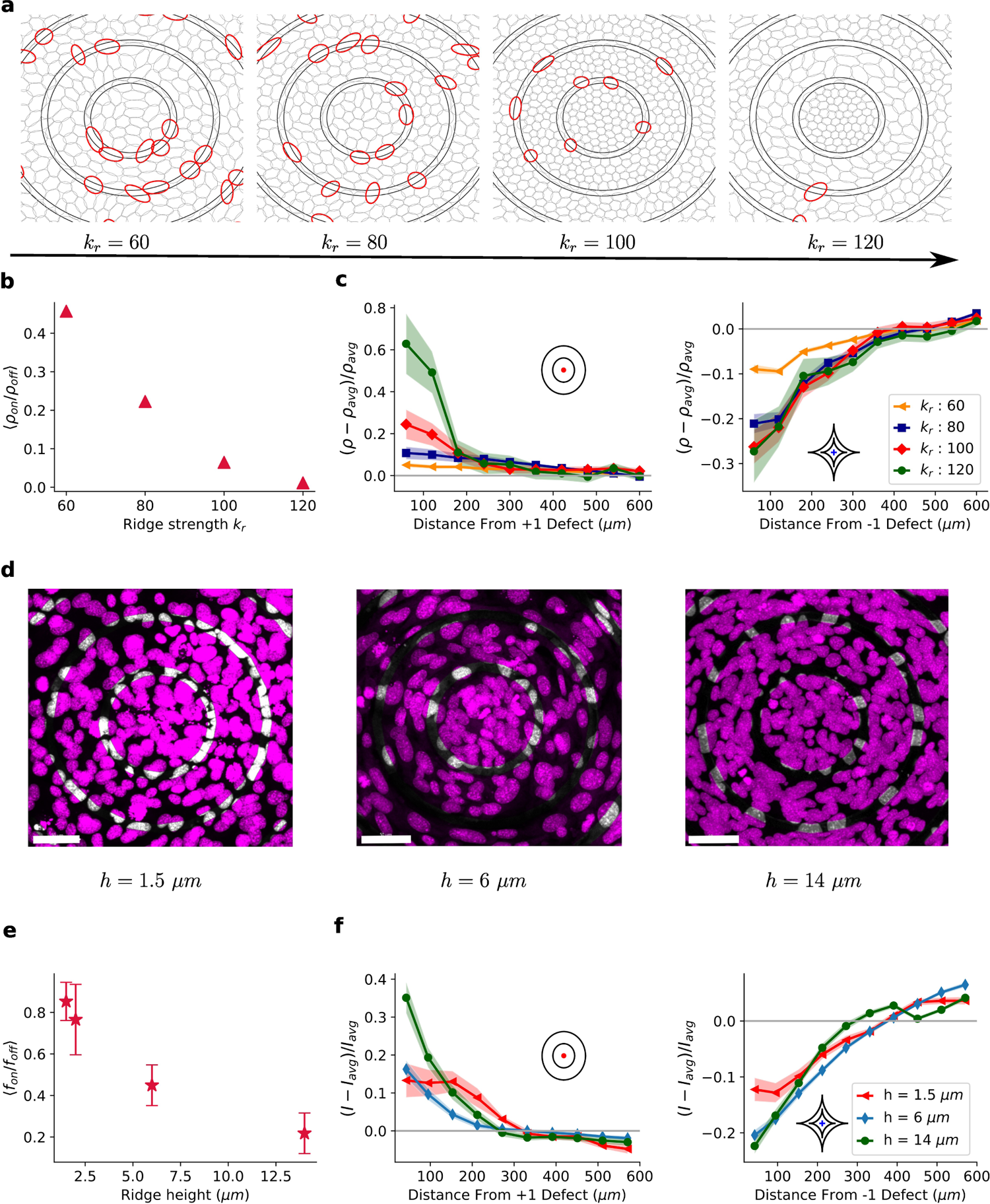
Ridge strength dependence of density variations. **a**, Snapshots of cells in simulations overlapping with ridges marked in red (non-overlapping shown in gray) for different values of ridge strength. A cell is considered to be overlapping with the ridges if the center of the ellipse is on the ridge. **b**, Ratio of number density of cells on and off the ridges averaged over 100 simulations. Error bars are smaller than marker size. **c** Density profiles of +1 (left) and −1 (right) defects for different ridge strengths. **d**, Confocal microscopy images (maximum intensity projections) of cell nuclei stained with Hoescht 33342. Parts of nuclei overlapping with ridges are shown in white and non-overlapping regions are shown in pink. **e**, Experimentally observed ratio of area fraction of cell nuclei on the ridges to area fraction of nuclei off the ridges for different ridge heights, determined from maximum intensity projection. **f**, Experimental density profiles at +1 (left) and −1 (right) defects for different ridge heights. The 1.5 *μm* data is from Fig. 1.

To experimentally test this prediction, we vary ridge height to constrain cell movements across ridges. Confocal microscopy of cell nuclei shows a reduction of fibroblast-ridge overlap as we increase ridge height from *h* = 1.5*μm* to *h* = 14*μm* (Fig. 6d). We quantify this by measuring the ratio *f*_on_/*f*_off_ between the fraction of the ridge area occupied by nuclei (*f*_off_) and the fraction of non-ridge area occupied by nuclei (*f*_off_). The average *f*_on_/*f*_off_ decreases roughly by a factor of four as the ridge height goes from *h* = 1.5*μm* to *h* = 14*μm*. Though *h* = 14*μm* is larger than a typical cell height, ridges of this height do not completely suppress crossing. We see that in these high ridges, the cell monolayer tends to become slightly undulated and three-dimensional, with cells not necessarily in contact with the substrate (Fig. S7). Effects of this three-dimensional structure will not be fully captured in our 2D simulations. Increased ridge height of *h* = 14*μm* leads to increased relative cell density near the +1 defect (Fig. 6d,f), consistent with our simulation results. Near the −1 defect densities remain low, with no clear systematic dependence on ridge height – similar to our simulation results for *k_r_* = 80 - 120. We see that by constraining cell migration and reducing ridge crossing, we increase density effects at the +1 defect core, as seen in our simulations, and as expected if cell division is driving the increase of density.

## III. DISCUSSION

We find that fibroblast alignment, velocity patterns, and density variation in our experiment are consistent with a model using shape dependent cell division. We experimentally induced defects using ridges, which resulted in high fibroblast density near +1 defects and low density at −1 defects. However, unlike experiments with other cell types where cell accumulation at positive defects and depletion at negative defects were driven by collective migration [9–11], fibroblasts did not manifest collective inward flow with highly aligned velocities. Instead, they moved in random azimuthal directions relative to the center of the +1 defect. This is consistent with earlier work arguing fibroblast monolayers are less driven by migration and activity [25]. To understand why density differences arise, we modeled fibroblasts as deformable elliptical cells. Our simulations found patterns of cell migration and alignment with the ridges that are consistent with experimental observations. Based on prior experiments on dependence of cell cycle progression on shape, we proposed a proliferation procedure where larger isotropic cells have higher probability to divide. This mechanism leads to density variations consistent with those in experiments. We predicted, using our simulations, that restricting cell movement across ridges would increase density at the +1 core – and confirmed it in experiments by modifying ridge height. Despite strong migration constraints, the marked density differences at defects were still present, which implies that cell division is important for explaining accumulation of cells.

Our model assumption that cell shape and size regulate division is consistent with past experiments [22, 23]. However, other experiments have argued that stress or pressure control proliferation and growth [26–30]. These may be elements of a single core mechanism, as cell shape, size and stress are all intertwined [31].

Our results imply that patterned substrates can regulate development of tissues via control of proliferation, not merely through controlling migration [32–35]. Similar approaches may help use mechanical cues to organize cells with limited motility. Beyond simply growth, other work shows that confinement and topology can provide cues to drive differentiation of cells [36, 37]. As cell area and aspect ratio are known to be important in determining the fate of individual cells [38, 39], our work suggests that ridge patterns could be harnessed for controlled development [40, 41] – but the observed feedback between growth and cell shape means that computational modeling will be required to understand the effect of any given pattern. Changes of cell shape and aspect ratio are also seen in many patterning processeses in development, including avian skin morphogenesis [42–44]; control of division by local cell shape may allow for additional feedback between tissue growth and local alignment. Our results suggest capturing the interplay of division, liquid-crystal alignment, and cell shape together is required to understand many patterning processes in eukaryotic development.

## IV. METHODS

### A. Simulation

We model cells as self-propelled elliptical particles with area *A* = *πab* and aspect ratio *AR* = *a/b* where *a* and *b* are major and minor axis radii respectively. The *a* and *b* can vary from cell to cell and will change over time; in our convention *a* is chosen such that it is the larger axis of the cell, *a* ≥ *b*. We perform Monte Carlo simulations using the Metropolis method. Briefly, we propose changes to cell properties – these changes are accepted with probability min(1, *e*^−Δ*E/T*^) where Δ*E* is the change in energy due to the proposed move, and *T* a temperature. As in, e.g. the Cellular Potts Model and related models [45], this is not a physical temperature, but a value setting the likelihood of fluctuations of different sorts. We choose the temperature *T* = 1 to set the energy scale of the problem. The three central elements of simulation are proposed moves, associated energies of the move, and cell division.

#### 1. Proposed moves

In one Monte-Carlo Step (MCS) we iterate over all cells in random order and propose a single move for each of them. For a cell *i* one of the four possible moves is attempted:

1. Move by 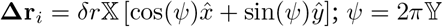.
2. Rotate by 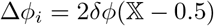.
3. Change major axis radius by 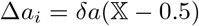.
4. Change minor axis radius by 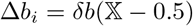.

where 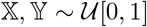 are random variables sampled from the uniform distribution defined in unit interval [0,1). The parameters *δr, δϕ, δa* and *δb* represent the maximum possible displacement, rotation angle, change in semi-major axis length and change in semi-minor axis length at each attempt, respectively (numeric values are given on Table I). We reject or accept a move after each attempt based on energy change Δ*E_i_* of the cell i due to the proposed move.

**TABLE I.**
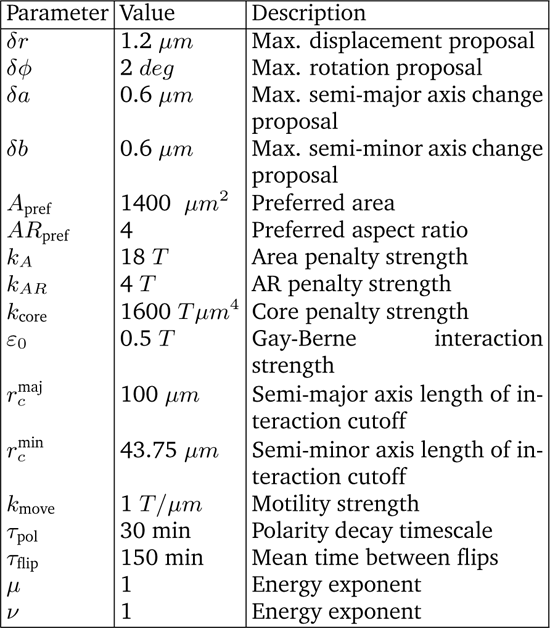
Default numeric values for parameters used in simulations.

To calibrate timescales of cell growth and speed, each of the proposed move types has a different probability to be selected. We propose displacement with probability of 10%, rotation with 20% and the two axis length changes each have 35% probability to be selected as a move. We have chosen this in part to ensure that cells quickly reach their steady-state shape, reflecting observations in experiments that, e.g. equilibration of fibroblast shape after division is much faster than significant motility [46].

#### 2. Cell Energies

We accept moves following the Metropolis criterion, which depends on the change in energy from a move. The total energy of our system is comprised of four distinct parts: geometric energies, cell-cell interaction energy, cell-ridge interaction energy, and motility energy. We describe each of these here.

##### Geometric energies penalize deviations from preferred size and shape

Cells have preferred area *A*_pref_ and preferred aspect ratio *AR*_pref_. We penalize deviations from preferred values with energy cost quartic in relative deviations *δ_A_* = (*A* – *A*_pref_)/*A*_pref_ and *δ_AR_* = (*AR* – *AR*_pref_)/*AR*_pref_:

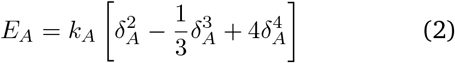

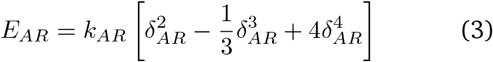

where *k_A_* and *k_AR_* are area and aspect ratio penalty strengths. The shapes of energy curves are shown in Fig. S8. Our goal in choosing these functions is to allow cells to easily change area and aspect ratio over a range of values close to their preferred values without significant energy cost. This reflects, e.g. for the area, that the cell can expand its height above the substrate, allowing it to make small area changes relatively easily. However, larger deviations result in substantial energy cost.

The finite size of organelles and high nucleus stiffness relative to the cytosol implies that cells can not be squeezed indefinitely. We model this feature via a core energy that introduces high energy cost if cell gets tiny, but is much smaller when cells have a typical size,

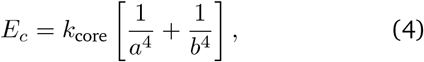

where a and b are the major and minor axis radii. We plot this curve in Fig. S8.

##### Cell-cell interaction energy favors parallel alignment of long axes of cells

Cells interact with neighbors within cutoff region via a modified Gay-Berne potential that is extensively used in liquid crystal simulations. The potential favors mutual alignment of cells and it is strongly repulsive when cells are too close and weakly attractive if cells are separated by longer distances.

Here we provide brief overview of the potential that we adapted for our simulations, detailed information can be found in [13, 47]. The interaction depends on relative orientations of cells. We characterize orientation of a cell *i* at position ***r**_i_* = (*x, y*) by unit vector ***û*** = (*u_x,i_, u_y,i_*) = (cos *ϕ_i_*, sin *ϕ_i_*), where *ϕ_i_* is the angle the major axis of the cell makes with the x-axis of simulation box. The interaction energy of a pair of cells located at positions ***r***_1_ and ***r***_2_ is given by:

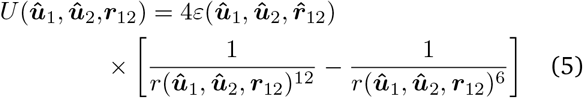

where ***û***_1_, ***û***_2_ are cell orientations and ***r***_12_ = ***r***_1_ – ***r***_2_ is a vectorial distance between centers of cells (Fig. S9). The function *r*(***û***_1_, ***û***_2_, ***r***_12_) is a scaled and shifted distance between cells:

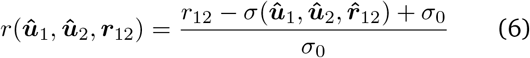

where 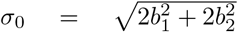 and 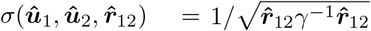 is an anisotropic range parameter that depends on size and orientations of cells via matrix *γ* that depends on the size and orientation of both cells 1 and 2 as:

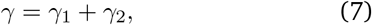

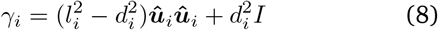

where 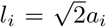 and 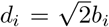 and the *I* is an identity matrix, and ***ûû*** indicates the dyadic product.

The term 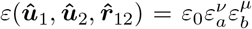 is an anisotropic interaction strength. Here, *ε*_0_ sets the general strength of interaction while 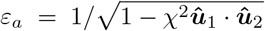 and 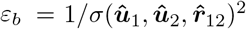 scale strength based on size and relative orientation of cells. The *ν* and *μ* are adjustable exponents set to 1 in our simulations and *χ* is given by:

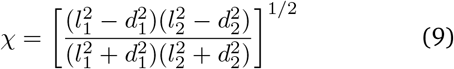

While the Gay-Berne potential of Eq. 5 has a longdistance attraction, we cut it off after a characteristic distance, reflecting that we do not expect cells to interact too far beyond contact. We compute the interaction energy of cell *i* with cells whose center is located within an elliptic area surrounding *i*. The elliptic area has same orientation as cell *i* and has semi-major and semi-minor axis lengths 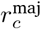 and 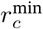 respectively (Fig. S9). This cutoff also allows us to speed up our simulations, as we only need to compute pairwise interaction energy between cells that are within this distance, which we track with a neighbor list. We update neighbor lists for all cells every time a cell divides or if any one of the cells moves by more than 25 *μm* with respect to its location during previous neighbor list update.

##### Ridge-cell interaction energy

Ridges are elevated with respect to rest of the substrate, so we penalize cell-ridge overlap. The energy cost of overlap is equal to the product of the ridge strength *k_r_* and fraction of the cell intersecting with the ridge, which we call ϒ:

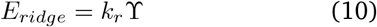

To estimate the fraction of cell overlap ϒ we compute the overlap between individual points within the cell, where these points are sitting on three ellipses with same orientation and shape of the cell. The axis lengths of the outermost ellipse match the cell size (*a, b*), and the inner two ellipses have axis radii (2*a*/3, 2*b*/3) and (*a*/3, *b*/3) respectively. If the number of points on these three ellipses that overlap with ridges is *N_o_* and total number of points is *N_t_*, then the fraction is given by ϒ = *N_o_/N_t_*. Each “feeler” ellipse has 64 points separated evenly in polar angle (Fig. S9).

##### Cell motility energy promotes movements along long axis in the direction of polarity

Cells are animate entities that constantly convert chemical energy into mechanical movement. They often have persistent direction of motion that may change by itself or due to cues like electric field, chemical gradient etc. [17]. The direction in which cell wants to travel is called (migrational) polarity, it points from rear of a cell where myosin contractions pull the “back” of the cell to the “front” where filopodia or lamellipodia push the frontier of the cell [48]. We denote the polarity vector of a cell by ***p*** = (*p_x_, p_y_*). In our model, when a cell rotates by Δ*ϕ* we correspondingly rotate the polarity. Because fibroblasts tend to move along the long axis of the cell [49], we choose the energy to promote motion along the long axis in the direction of polarity. For instance, if **Δ*r*** = (Δ*x*, Δ*y*) is a proposed displacement of one cell with polarity **p** then the motility energy change that results from this move is:

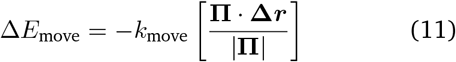

where **Π** = (***û*** · ***p***)***û*** is projection of polarity onto the long axis of the cell. Note that, in our approach, the magnitude of the polarity is irrelevant – only its direction contributes to the energy. *k*_move_ sets the relevance of the motility compared to other driving forces. The energy we use here is akin to, e.g. energies used in the Cellular Potts Model to represent cell polarity and motility [21, 50].

The polarity has positive feedback with the displacement in one MCS, **Δ*r***, which could also be zero if no displacement is proposed or proposed displacement is rejected. At every MCS t, we update polarity as in Szabo et. al. [21]:

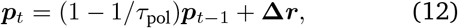

where ***p**_t_* and ***p***_*t*–1_ denote polarity new polarity at step *t* and polarity at previous step *t* – 1 respectively, and the *τ*_pol_ is a polarity decay timescale parameter measured in units of MCS. If there was no displacement Δ*r* = 0, polarity would decay in magnitude. However, because displacements tend to correlate with polarity, and correlate more with polarity if many cells are locally pushing in the same direction, Eq. 12 tends to cause cells to locally align. The effect of changing polarity decay time has been systematically investigated by [19]. In this sort of model, increasing the time required for polarity to decay ensures that the polarity is largely controlled by the sum of previous displacements over a long time – generally making the migration more coherent.

In addition, since fibroblasts periodically reverse direction of motion [4], we stochastically flip cell polarity, **p** → −**p**. Reversal happens with probability 0.01 every 1.5 minutes (100 MCS). The number of tries needed for flip event has geometric distribution with success probability *p* = 0.01. Expected number of attempts needed for reversal is then 1/*p* = 100 (i.e. average flip time is *τ_flip_* = 1.5 × 100 = 150 minutes). This random flipping disrupts polar coherence, preventing cells from forming a uniformly rotating state as can be seen in, e.g. experiments on epithelial trains [51].

#### 3. Cell division

As in the monolayer experiments, in simulations our cells divide and proliferate. We initialize our system by putting circular cells of initial radius *r*_0_ = 10 *μm* at random positions (excluding configurations with cell-cell overlap) at a density of *ρ* ~ 70 cells/mm^2^. We let cells evolve for 10h without division, to ensure they can relax to reasonable shapes. Then we divide one cell every 1.5 minutes (100 MCS), choosing this rate to roughly match the experimental growth curve (Fig. S1), and halt division once cells reach their terminal density of *ρ_f_* ~ 2000 *cells/mm*^2^ (3000 cells in our simulation box size of 1200 *μm* × 1200 *μm*). In our model, the number of cells as a function of time is always the same from simulation to simulation, but which cell divides at any point is stochastic. Cells have shape dependent probability to be selected to divide: given *N* cells in the simulation, cell *i* is selected to divide with probability

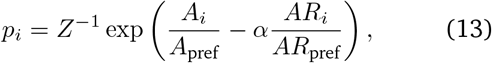

where *α* is parameter that tunes sensitivity of probability to cell shape and *Z* is a normalization factor chosen such that 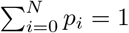.

When cell *i* of size (*a_i_, b_i_*) and orientation *û_i_* divides the two daughter cells both will have size (*a*_1_, *b*_1_) = (*a*_2_, *b*_2_) = (*a_i_*/2, *b_i_*/2) and orientation *û*_1_ = *û*_2_ = *û_i_* (Fig. S9). Divisions occur along the long axis of the cell; the choice of daughter cell sizes ensures that the total length of the cell is preserved. The choice of daughter cell size does mean that area is not conserved in the division – this is in part to avoid potential numerical problems with extreme cell-cell overlap which can be caused by division. We find that cells quickly grow up to a size comparable to nearby cells post-division when possible.

#### 4. Randomness and seeds

When varying parameters (e.g. in Fig. 4, Fig. 6), we perform 100 simulations (indexed 1, 2,…, 100) for each parameter set. To better understand the effect of the changed parameter, we keep the random number generator seed of each simulation fixed – so there are 100 distinct seeds, and when a parameter is varied, e.g. comparing *α* = 2 to *α* = 0 in Fig. 4, we are comparing simulations that have the same initial conditions and seeds. This choice is made to make it clearer that changes are systematically due to the effect of the parameter change, and not randomness. However, because the initial conditions are the same for all of our runs, we need to be confident that these initial conditions are not significantly driving our results. We provide a test of this in Fig. S10, swapping initial conditions between the +1 and −1 defects. While there is some quantitative difference from earlier results in Fig. 4, we still see the key results that density is increased at the +1 defect as we make *α* → 2.

#### 5. Broader modeling considerations

One contribution of our work is a new cell-based framework, which can describe collective migration of highly anisotropic cells while resolving individual cell shapes and positions. This is one of many possible approaches to modeling collective cell migration [52, 53]. We discuss some of the broader choices we made here. We argue continuum tissue / active liquid crystal models [53, 54] would be inappropriate to model these experiments, as continuum models are restricted to length scales that can average over many individual elements – incompatible with studying cells of typical width *~* 20 microns in ridges with spacing 60 microns. Our approach also avoids issues with orientational anisotropy associated with lattice models like the Cellular Potts Model [45]. In principle, phase field approaches [20, 55–59] would also avoid lattice artifacts, but our scale of ~ 3000 cells is an order of magnitude larger than typical applications of even simplified phase-field models [60–62]. Earlier papers have modeled elongated self-propelled objects with particle-field and/or Gay-Berne approaches [63, 64], though without explicitly describing deformability. Our approach is probably closest to the deformable self-propelled particle approach of Menzel and Ohta [65] and the deformable ellipsoids of Palsson and Othmer [66].

We have neglected in our paper the possibility that cells may create alignment of fibronectin or other extracellular matrix proteins that may play a role in long-range guidance [67]. We were able to recapitulate alignment to ridges without this effect. However, it may be essential to understand longer-scale perfect alignment on unpatterned substrates [4]. We also neglect potential couplings between cell shape and polarity [68–70], which can drive complex behaviors like cell circling and oscillation in response to fields, as observed in keratocytes [69]. We have neglected these factors because we have no evidence that fibroblasts show circular migration behavior similar to keratocytes.

We have focused on how our division rules alter local effects of density in response to patterning. These division rules will also likely affect the mechanical properties of the tissue and the degree of fluidity [71, 72]. These are factors that might be important to study further in extensions to more motile cells than our 3T6 fibroblasts.

### B. Experimental Methods

#### 1. Cell culture

The 3T6 mouse fibroblasts (from ATCC) are cultured in CellTreat tissue culture dishes using 90% Dulbecco’s Modified Eagle’s Medium (DMEM) [+] 4.5g/L glucose, L-glutamine, sodium pyruvate (Corning CellGro) and 10% Fetal Bovine Serum (Corning CellGro). When outside the incubator for long-duration (>30 minutes) imaging, the growth medium is replaced with 90% CO**2** Independent Medium (Gibco) and 10% Fetal Bovine Serum (Corning CellGro), with 4.5 g/L L-glutamine added (Quality Biological). Cells are utilized only up to generation 20.

##### Growth curve

The experimental growth curve of Fig. S1 is obtained using a standard method of seeding cells onto 5 Petri dishes with a density of 60 cells/mm^2^. To measure this seeding density, a subset of the suspended cells are stained using Trypan Blue and counted using a hemacytometer (10*μL* volume). Every day, one dish is selected, for which cells are resuspended and counted with the same method. Cells in the 5 dishes are counted after 24, 48, 75, 95, and 120 hours respectively.

#### 2. Substrate preparation

The topographic features are patterned using a mold of SU-8 on glass. SU-8 is a negative photoresist, a hard polymer which crosslinks by exposure to UV light. To create 1.5*μ*m-tall ridges, we use SU-8 2002 (MicroChem), by spincoating the SU-8 at a maximum speed of 3000rpm for 30 s. To create 6*μm*-tall ridges, we use SU-8 2005 (MicroChem), by spincoating the SU-8 at a maximum speed of 3000rpm for 30 s. To create 14*μm*-tall ridges, we use SU-8 2010 (MicroChem). In this case, we use a maximum speed of 2000rpm for 60s.

Polydimethylsiloxane (PDMS) substrates are prepared from Sylgard 184 (Dow Corning) with 15% curing agent. After mixing, the PDMS is degassed at room temperature and then poured over the SU-8 patterned substrates. These are cured in a vacuum oven at 60°*C* for 2 hours. The patterned PDMS is then prepared for cell culture.

We submerge the patterned PDMS in ethanol for 20 minutes to sterilize it. The substrates are then dried at room temperature, and then coated with a minimal volume of fibronectin from bovine plasma (MilliporeSigma, 25*μm*/*mL* in PBS) for 45 minutes at room temperature prior to use for cell culture.

Prior to seeding onto the patterns, a subset of the suspended cells are stained using Trypan Blue and counted using a hemacytometer (10*μL* volume). Cells are seeded at a concentration of 5 × 10^5^ cells/dish, with each dish being circular with a diameter of 100mm. This corresponds to 60 cells/mm^2^.

We note here that the data for cells growing on 1.5*μ*m shown in Figure 1d and in Figure 6f are from our previous publication [12], For these data sets, the seeding density was not held constant.

As a control, the ridge height and pattern quality of the PDMS with ridges up to 6 *μ*m are verified using a Keyence VK-X200K color 3D laser scanning microscope. The 14*μ*m ridge heights are verified by imaging the crosssection of the PDMS using a 50X objective on a Nikon LV Pol microscope with a Nikon DS-Ri2 camera. For all ridge heights, we analyze the images in the acquisition software by measuring the height of the top surface at a series of locations both on and off the ridges and computing the average difference, with standard error as uncertainty. Measurements of ridge height are shown in Fig. S11.

#### 3. Cell imaging

##### Preparation/Staining

The cells’ nuclei are stained using Hoechst 33342 dye (10 mg/mL stock solution, Invitrogen), diluted at a ratio of 1:1000, followed by 15 minutes in the incubator. When they are stained for the purpose of acquiring a video, the dye solution is removed, and the dish is filled with CO**2**-independent media.

For the samples with 1.5*μ*m high ridges, there are 3 images each of cells around +1 defects and −1 defect of which the nuclei are stained using NucRed Live 647 (Invitrogen) rather than Hoechst 33342. The dye is added following the suggested protocol of 2 drops/mL of media, followed by 15 minutes of incubation. Data about nuclear orientation and cell density from these images are incorporated into Figure 1c, Figure 1d, and Figure 4b.

For confocal images, cells are fixed before being stained with Hoechst 33342 and Phalloidin-iFluor 594 conjugate (AAT Bioquest). For fixing, the cells are first incubated in 4% paraformaldehyde for 10 minutes at room temperature. Then they are washed with PBS three times, each time incubated at room temperature for 5 minutes. Then, they are incubated in PBS with 0.1% Tween-20 at room temperature for 10 minutes, followed again by washing three times in PBS for five minutes each. Once the cells are fixed, they are stained with a solution with 10*μ*g/mL Hoechst 33342 and a working solution of the Phalloidin stain. This Phalloidin stain is comprised of 1 *μ*L of Phalloidin-iFluor 594 Conjugate solution (AAT Bioquest) diluted in 1 mL of PBS with 1% Bovine Serum Albumin. The cells are incubated in this solution for 20 minutes prior to imaging.

##### Microscopy

Phase contrast and fluorescent imaging in 2D is done using a Nikon TI-Eclipse microscope using a Hamamatsu Orca-flash camera. Large format images are acquired by translating the stage with 15% overlap between adjacent frames, and then the images are stitched together using Stitching (Grid/Collection stitching) plugin in ImageJ [73]. At each location, we take one phase contrast and one fluorescent image.

For video acquisition, phase contrast and images of nuclear fluorescence are taken at 6 minute intervals. While acquiring the video, the Tokai Hit ThermoPlate microscope stage is heated to 37°*C*, and the stage is covered with a plastic sheet. Twice a day, or as needed, fresh CO_2^-^_ independent media is added to refill the dish, which loses medium due to evaporation. Autofocus is performed in NIS-Elements (Version 5.02.01) before each image is captured, and the light is switched off between frames.

##### Confocal microscopy

To quantify the prevalence of cells growing over ridges, the cells are imaged using a Leica SP8 confocal microscope with a White Light Laser and Leica HyD detectors and 40X objective. They are imaged after being fixed, permeabilized, and stained with Hoechst 33342 dye (10 mg/mL stock solution purchased from Invitrogen), diluted at a ratio of 1:1000. For each step, we take a stack of images at different heights from 3 channels: a bright field channel, a channel collecting the information from the nuclear fluorescence, and a channel collecting the information from the actin filaments stained with Phalloidin 594.

#### 4. Image analysis

##### Confocal microscopy

The images are segmented with the software IMARIS 9.8.2 using the “Create Surface” feature to identify the cells growing over the ridge from those growing between ridges. In the bright field images, we identify clearly defined lines on the bottom plane of the z-stack corresponding to the location of the edges of ridges. Using a drawing tool in IMARIS, we generate a shape on each edge. Then on the top plane of the z-stack, we paste the same shapes that were generated on the bottom surface. Around the circular ridges used to generate +1 defects, this demarcates a cylindrical shell which we identify as the “on wall” region of the image. This procedure is repeated for every ridge in every image. Once these regions are demarcated, we use the command “Mask Selection” to create a new color channel containing only voxels from the fluorescent image which are “on wall”. These regions, identified with the method described above, are used to create a mask on the corresponding images of fluorescent nuclei.

Once the masks are made, the 3D images are analyzed using ImageJ. First, all the channels are analyzed separately; then the images are collapsed to 2D using the maximal intensity of a voxel at a given xy-position. An intensity threshold is applied to create binary images, only showing the nuclei at given xy-positions. Fractions of black/white pixels on/off walls are computed to identify the area fractions on/off walls occupied by nuclei. From these, we report the average area fractions and the standard deviations.

##### Cell alignment

Quantification of cell orientation also followed the protocol of [12]. Briefly, we define the axis along which the topographic features orient the cells as a function of its azimuthal coordinate with respect to the defect, following the protocol described in [12]. For a + 1 defect, this is the coordinate itself, modulo 180 degrees. For a −1 defect, this is 180 degrees minus the coordinate in the first and second quadrant, and 360 minus the coordinate in the third and fourth quadrant. We then compare the deviation in the direction of the cell’s major axis from this axis, which is the deviation in the cell’s alignment from the expected or patterned angle. The orientation of the cell’s major axis is determined by fitting ellipses to nuclei identified in the fluorescent images in ImageJ, first by creating a binary mask of the image.

##### Cell density vs. distance from defect

We identify the center of the nearest patterned defect from the phase contrast images. For each pixel in the fluorescence image of the nuclei, we compute the distance from the nearest defect. We then compute the sum of the intensities and the number of pixels in each 60 micron ring out to a distance of 600 microns, and from these values we compute an average nuclear intensity in each ring. In each plot we report the average and the standard error, after normalizing by dividing by the average intensity measured within 600 microns from the defect center.

##### Cell tracking

Cell tracking is achieved using the TrackMate plugin in ImageJ [74]. The simple Linear Assignment Problem (LAP) tracker is used, which does not detect merging and splitting events. The trajectories of each cell are imported into Matlab for further analysis. Fig. 1g shows the direction of displacement of the cells in the first hour of the video. Only the cell paths that are continuously identified for every frame in the first hour are included in the image. This includes the majority but not all of the cells in the frame. Then, these cells are identified as moving clockwise or counterclockwise, and a vector with a fixed length is drawn on the first frame of the video. Its direction indicates the direction (but not the magnitude) of the net displacement of the cell in the first hour and its color indicates whether the cell moves clockwise or counterclockwise.

## ACKNOWLEDGMENTS

We acknowledge Matthew Pittman for help with confocal microscopy. KK and BAC acknowledge support from NSF grants PHY 1915491, DMR 1929467, and NIH R35 GM142847. FS is funded by Novo Nordisk Foundation Recruit grant NNF21OC0065453. This research project was conducted using computational resources at the Maryland Advanced Research Computing Center (MARCC). KDE acknowledges the support of the National Science Foundation Graduate Research Fellowship under Grant No. DGE-1746891.

## Supplemental Appendix

**Movie 1**: Cell movement around +1 defect. 8 hour experimental video from fluorescence microscopy showing nuclei stained with Hoechst 33342 which have been tracked using TrackMate in ImageJ. h=1.5*μ*m and ridge spacing is 60*μ*m.

**Movie 2**: Time evolution of simulation with centered +1 defect from 100 cells to 3000 cells in 93 hours. Cells are colored according to angle they make with x-axis as in Fig. 2.

**Movie 3**: Same trajectory as Movie 2, but recentered such that −1 defect is at the core.

**Movie 4**: Simulation showing mixing of cells when ridge strength *k_r_* is set to 60. At the start of simulation cells are colored according to their radial location. If a cell is initially within the first (innermost) ring, it is colored orange, cells between the first and second ring are colored black, etc. All descendants of cell inherit color from their parent (or equivalently root) cell. We can see that cells that origins at different locations eventually cross ridges and mix.

**Movie 5**: Same as Movie 4, but for the ridge strength *k_r_* = 120. In this case the overwhelming majority of cells do not cross ridges.

**FIG. S1.**
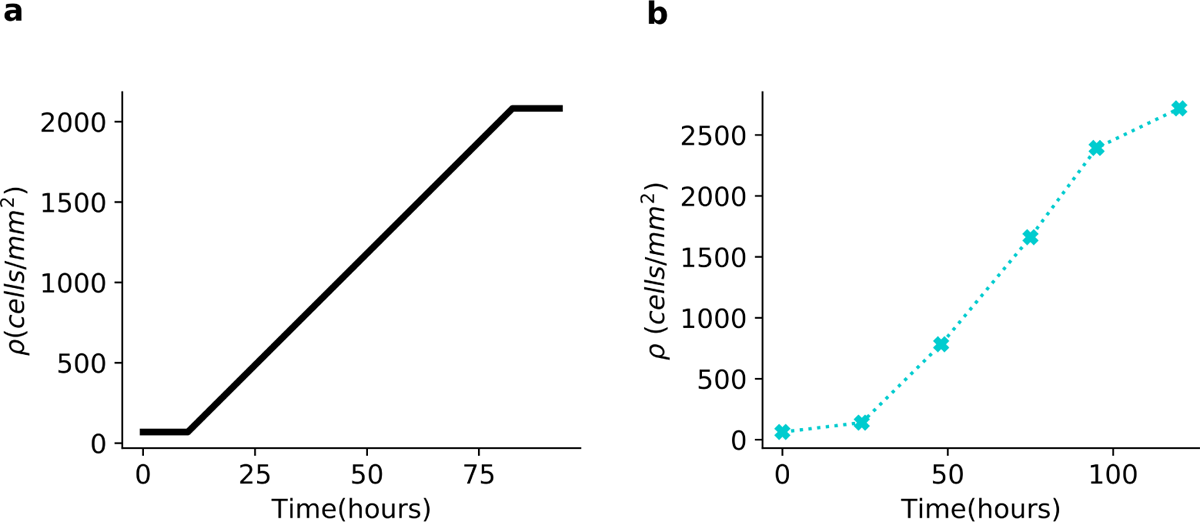
Growth curves show cell number increases over time and then begins to saturate. **a**, Simulation. Note this growth curve is imposed by our approach, and chosen to roughly match the experiment (see Methods). **b**, Experiment on 3T6 fibroblasts, conducted as detailed in Methods.

**FIG. S2.**
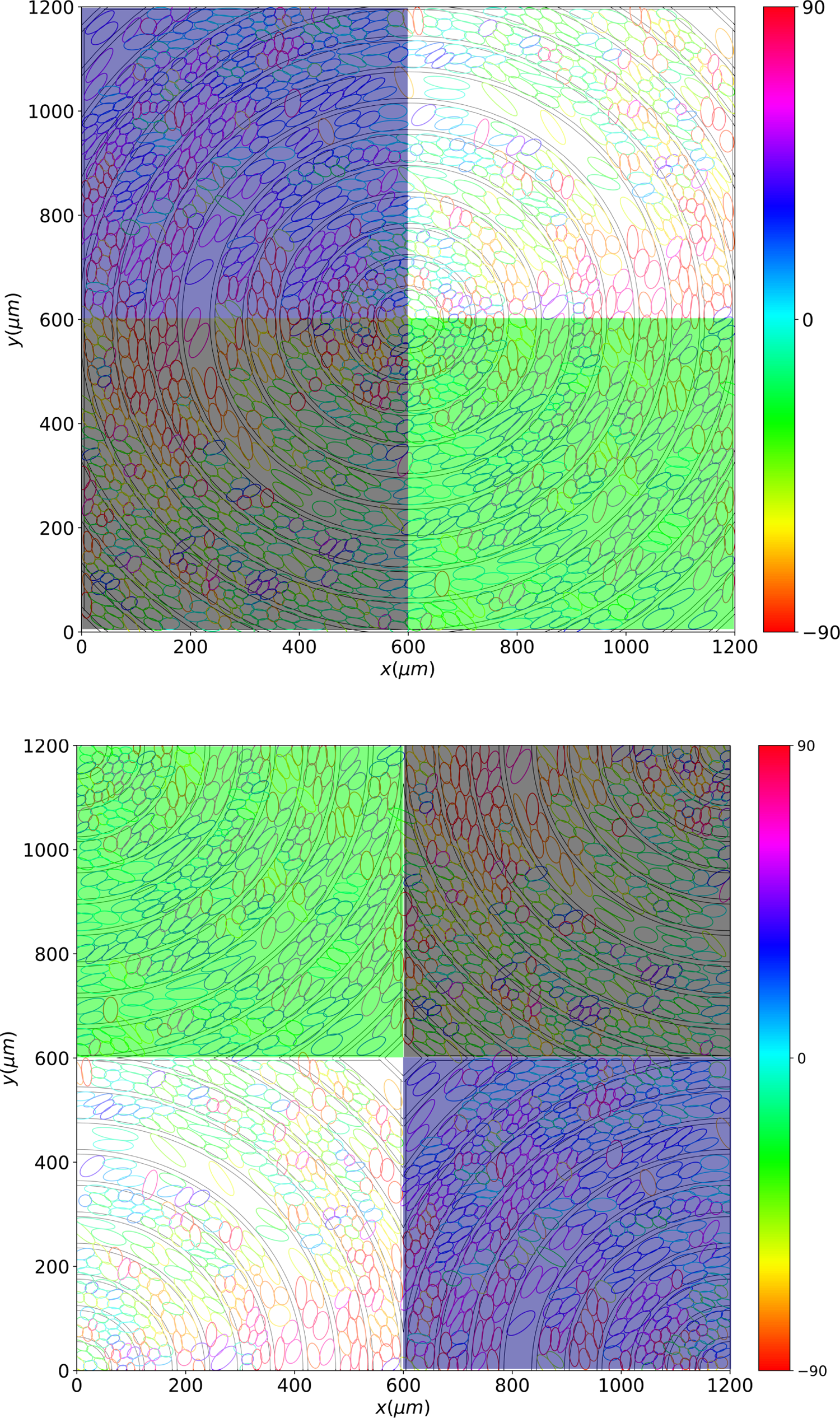
Simulation box has dimensions of 1200×1200 *μm*^2^ and periodic boundary conditions. If +1 defect is centered then −1 defect is at the corners of simulation box. The figure demonstrates of rearrangement of quadrants of the simulation box with centered +1 defect (top) that centralizes −1 defect (bottom). Color bars indicate the angle between major axis of cells and x-axis.

**FIG. S3.**
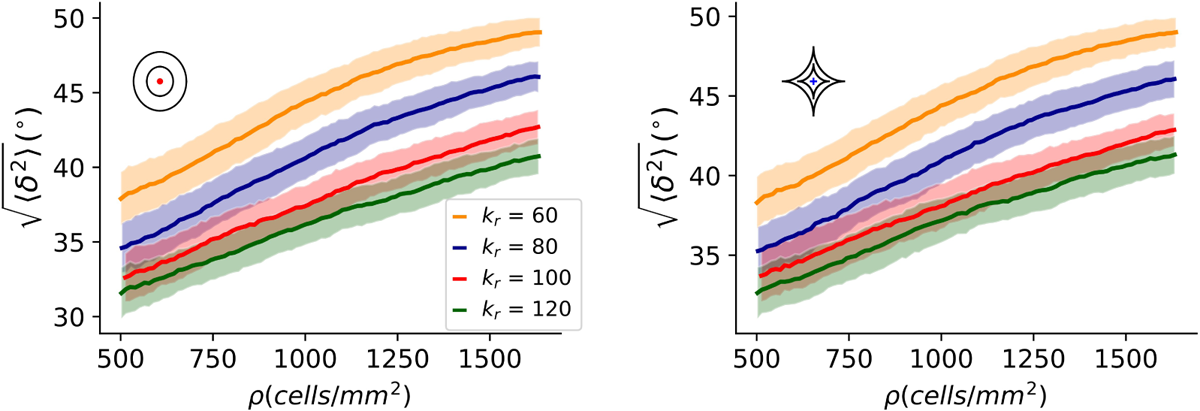
Simulation: RMSDs 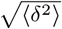 for different ridge strengths when cells are randomly selected to divide, i.e. *p_i_* = 1/*N*, and in the absence of motility energy (i.e. *k*_move_ = 0) for +1 (left) and −1 (right) defects.

**FIG. S4.**
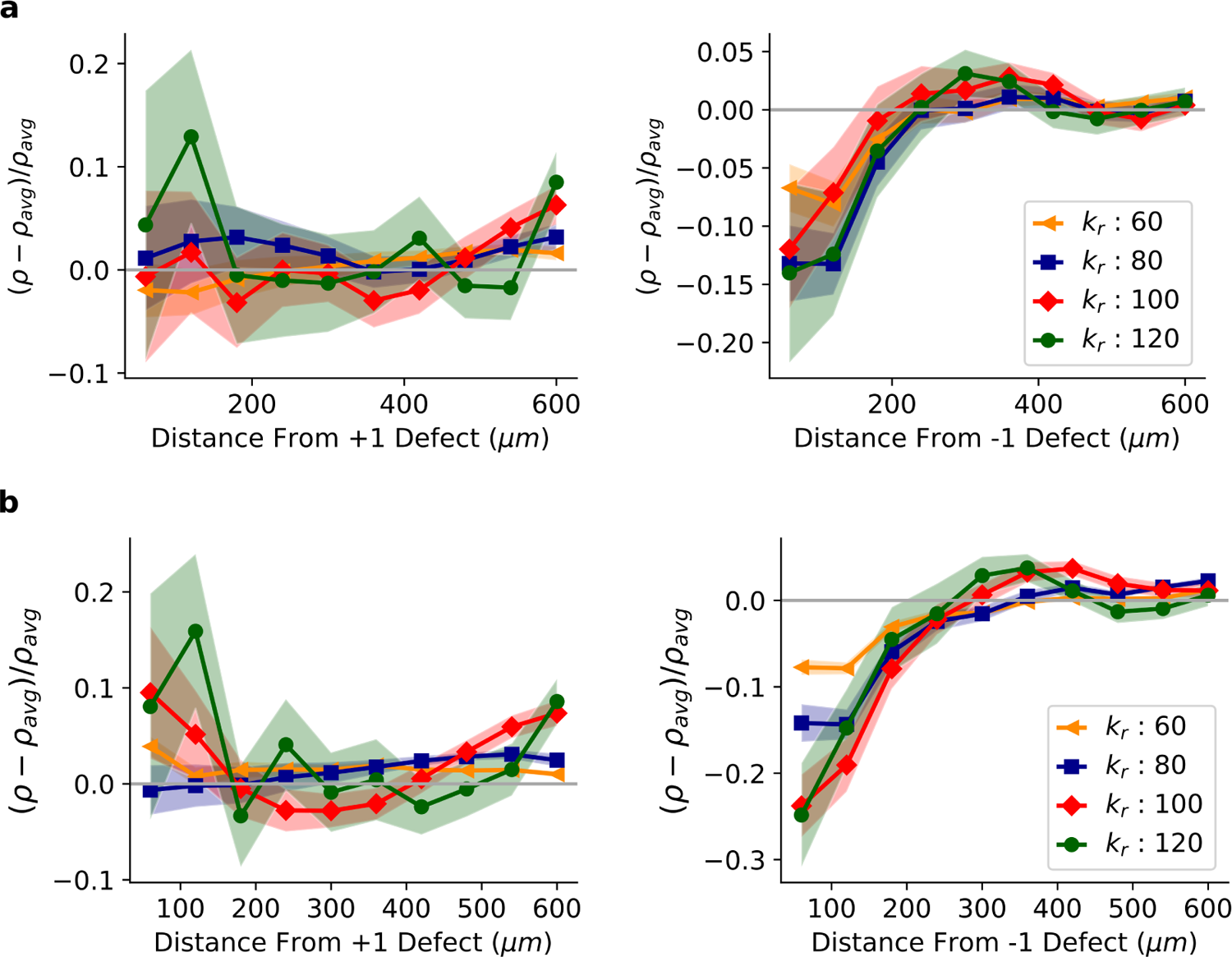
Simulation: Density profiles for different ridge strengths when cells are randomly selected to divide, i.e. *p_i_* = 1/*N*. **a**, Densities near +1 (left) and −1 (right) defects without motility energy (*k*_move_ = 0). **b**, Densities near +1 (left) and −1 (right) defects with motility energy (*k*_move_ = 1).

**FIG. S5.**
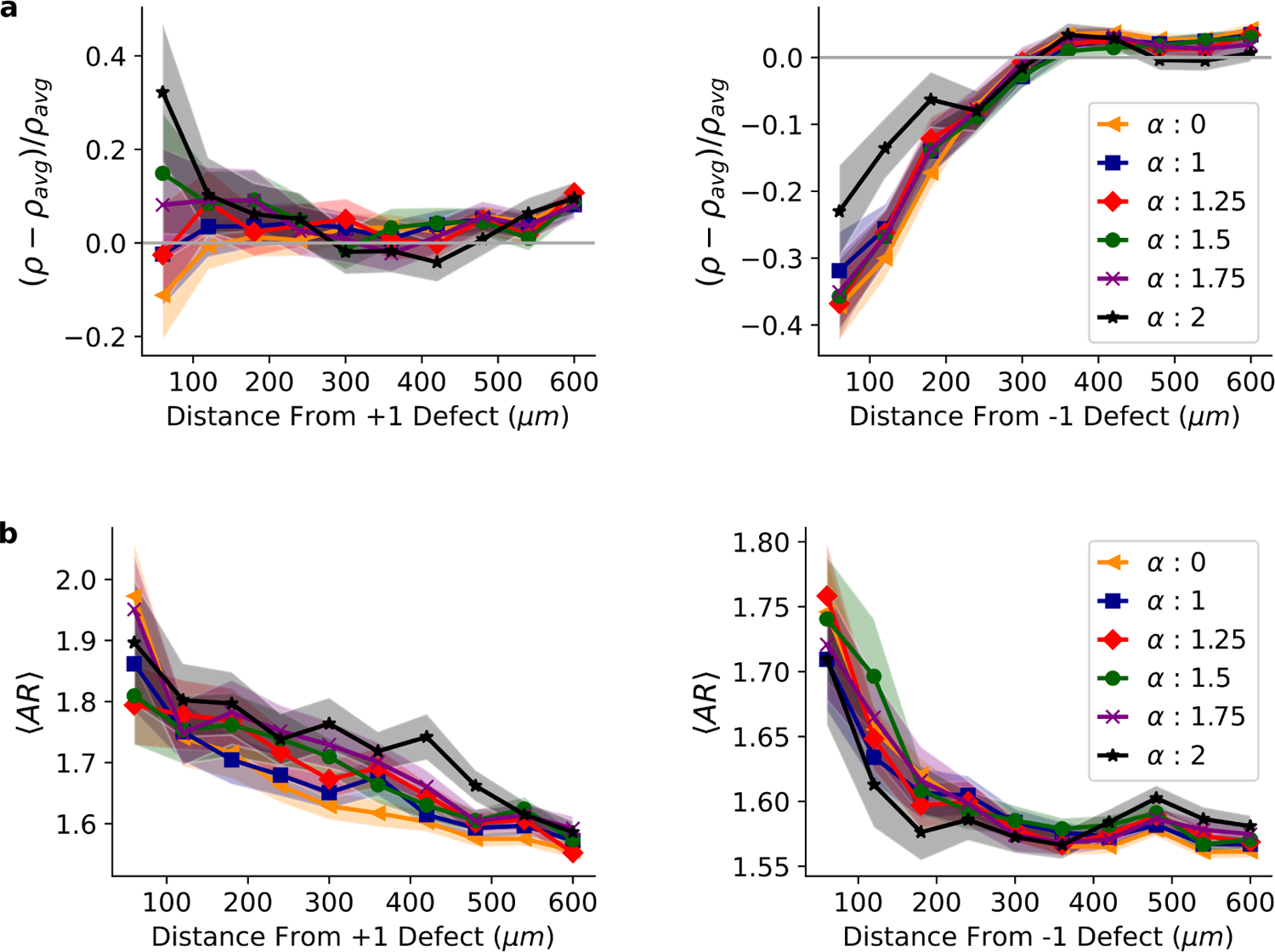
Simulation: Density and aspect ratio profiles in the absence of motility energy (*k*_move_ = 0) and ridge strength of *k_r_* = 120 for different sensitivity of division probability to shape *α*. **a**, Densities near positive (left) and negative (right) defects. **b**, Aspect ratios near positive (left) and negative (right) defects.

**FIG. S6.**
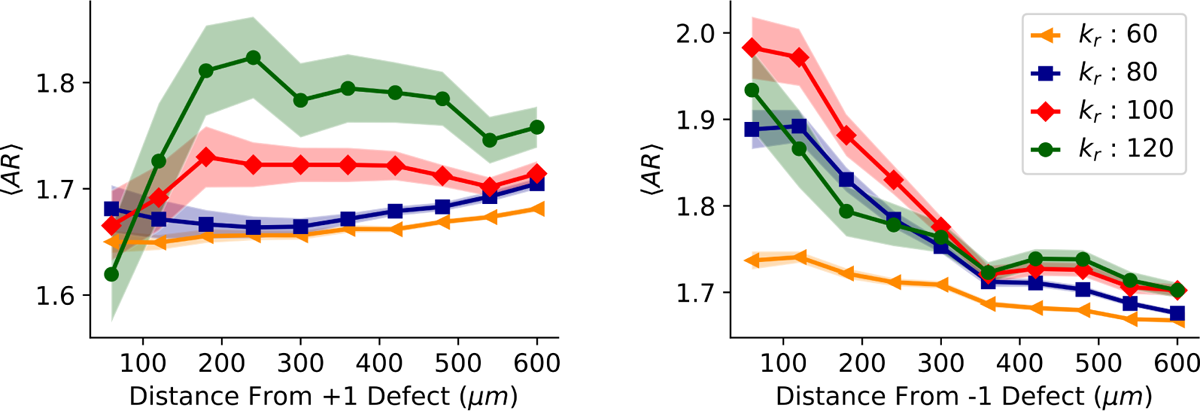
Simulation: Aspect ratio profiles for different ridge strengths near +1 (left) and −1 (right) defects. Here *α* = 2 and *k*_move_ = 1

**FIG. S7.**
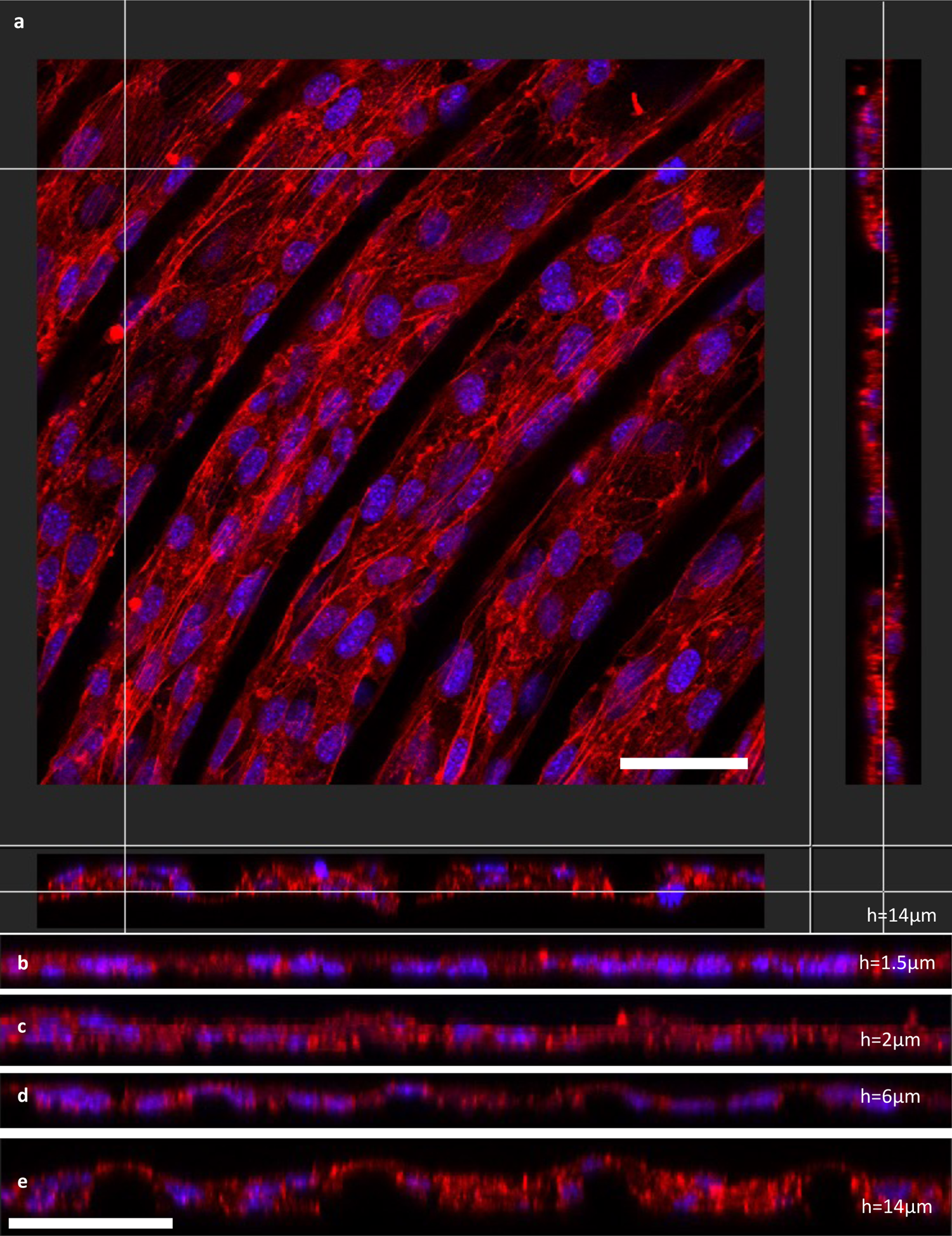
Experiment: Confocal imaging of cells to show 3D structure. **a**, Confocal image of nuclei (stained with Hoechst 33342) and actin filaments (stained with Phalloidin-iFluor 594 conjugate). This shows three orthogonal views, the top-down view and two side views, with the location of the cross-sections shown by the crosshairs. The scale bar is 50 *μ*m. **b**, Cross-section from a h=1.5*μ*m sample, showing the nuclei lying in a flat monolayer. Scale bar for **b-e** is 50 *μ*m. **c**,**d** Cross-section from h=2*μ*m and h=6*μ*msamples, respectively. As the ridge height increases, the monolayer becomes more disrupted, but the nuclei still occupy the same plane. **e**, Cross-section from a h=14*μ*m sample. In this case the monolayer becomes more undulating, and the nuclei can be seen at different heights, indicating that the cell environment is becoming more 3-dimensional.

**FIG. S8.**
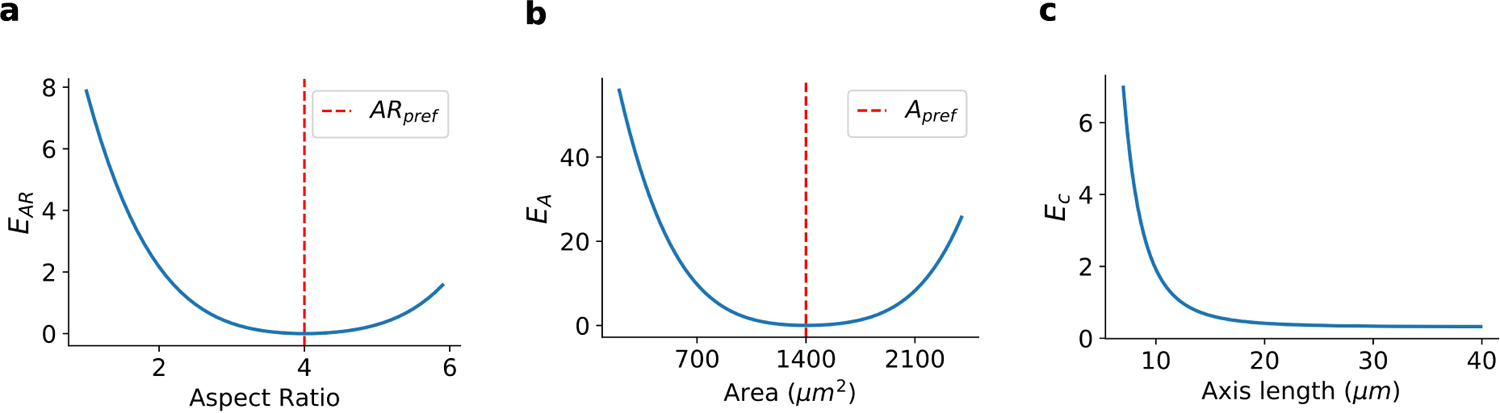
Geometric energies that penalize deviations from preferred shape and size. **a**, Quartic aspect ratio penalty for preferred aspect ratio *AR*_pref_ = 4. **b** Quartic area deviation penalty energy for preferred area *A*_pref_ = 1400 *μm*^2^. **c**, Core energy as a function of one of the axis lengths when the other axis is set to 15 *μm*.

**FIG. S9.**
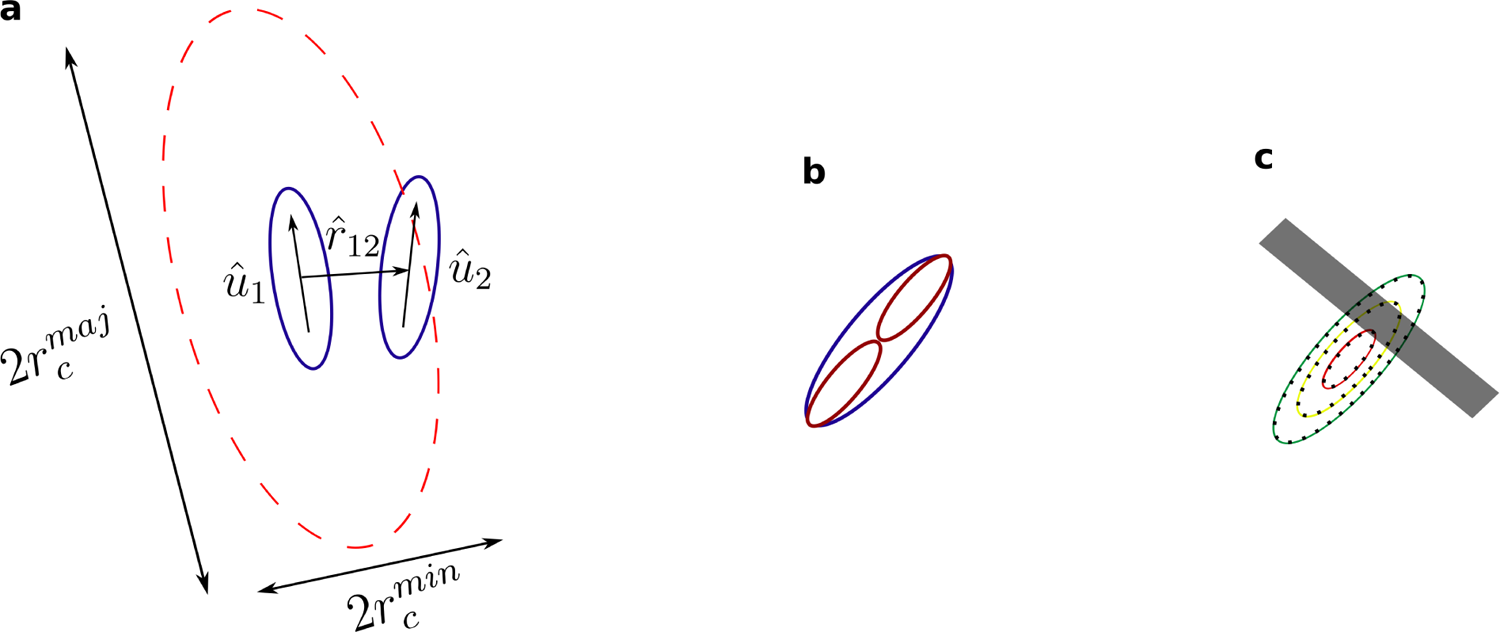
Illustrations for cell-cell interaction, division and cell-ridge interactions. **a**, Schematic of the Gay-Berne interaction cutoff region for cell 1 (with orientation *û*_1_) shown with red dashed ellipse. The ellipse has major axis length 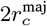 and minor axis length 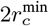. Cell 2 with orientation *û*_2_ is within interaction range. **b**, Illustration of a parent cell (blue) dividing into two daughter cells (red). **c**, Sketches of three ellipses with dimensions (a, b), (2a/3, 2b/3) and (a/3, b/3) that are used to sense ridge shown by gray area. The dark dots within shaded ridge area contribute to cell-ridge overlap penalty.

**FIG. S10.**
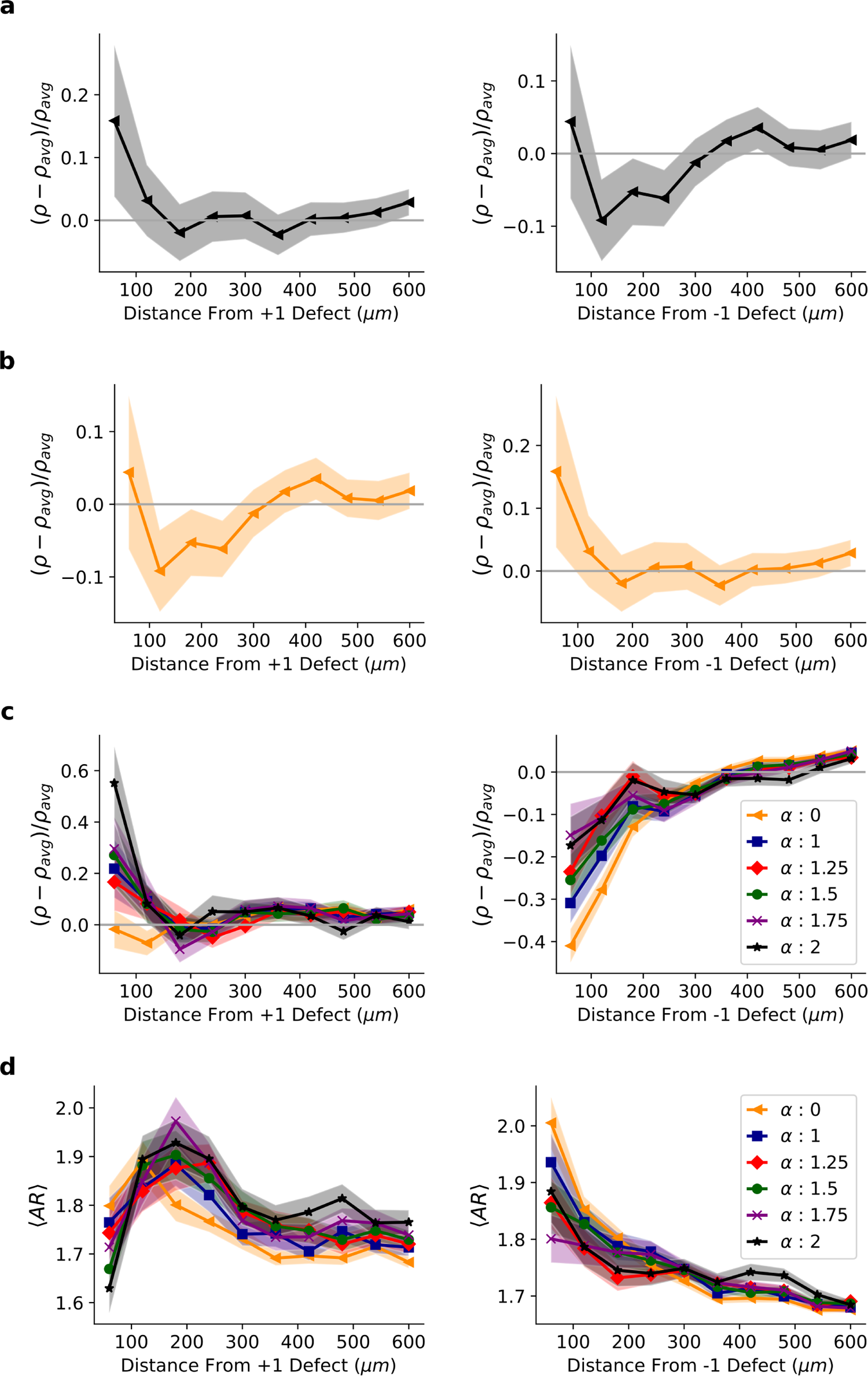
Simulation: Density and aspect ratio profiles when initial conditions are swapped. **a**, Default initial density profiles at the start of simulation for +1 defect (left) and −1 defect (right). **b**, Swapped initial conditions. End state densities (**c**) and aspect ratios (**d**) near +1 defect (left) and −1 defect (right) when initial conditions are swapped for different values of shape sensitivity α. Here *k_r_* = 120.

**FIG. S11.**
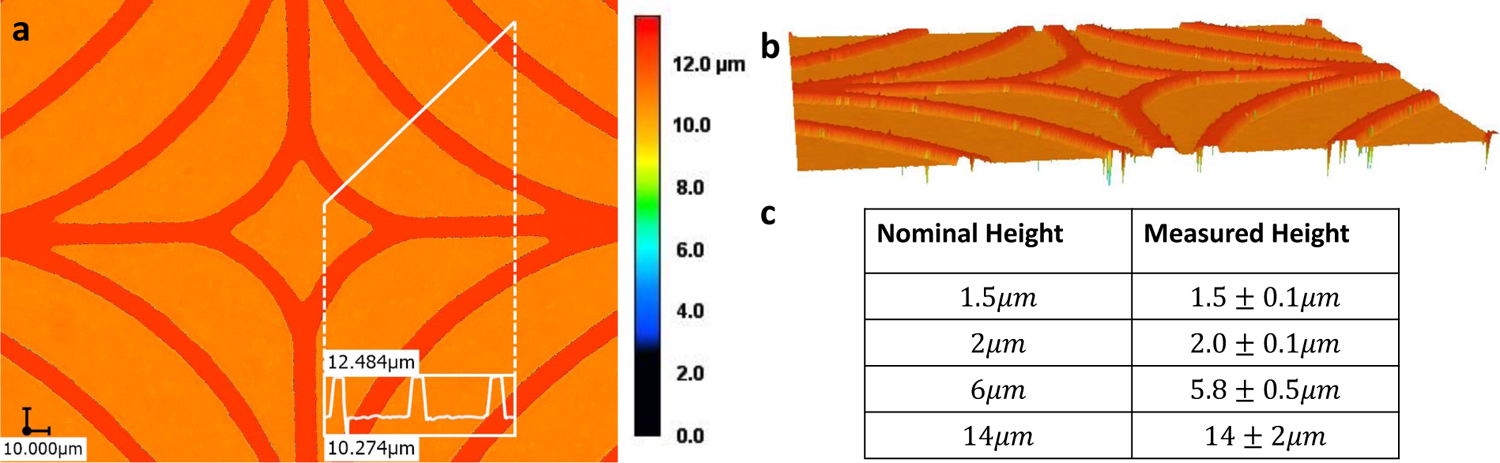
**a** Top-down view of h=1.5*μ*m PDMS acquired with the laser scanning microscope. The color represents height. **b** 3D rendering of the same image, with the colors also representing height. Here, the spikes below the surface at the edges of the ridges are artifacts resulting from scattered light. (c) Table of the measured heights (mean and standard deviation) of the PDMS samples. The ridge heights of the h=1.5*μ*m, 2*μ*m, and 6*μ*m samples were measured using the laser scanning microscope, and the h=14*μ*m sample was measured by viewing the cross section of the PDMS with the optical microscope with a 50X objective.

## References

[1] L. A. Davidson, Epithelial machines that shape the embryo, Trends in Cell Biology 22, 82 (2012).

[2] D. Andrienko, Introduction to liquid crystals, Journal of Molecular Liquids 267, 520 (2018).

[3] S. Shankar, A. Souslov, M. J. Bowick, M. C. Marchetti, and V. Vitelli, Topological active matter, Nature Reviews Physics 4, 380 (2022).

[4] G. Duclos, S. Garcia, H. Yevick, and P. Silberzan, Perfect nematic order in confined monolayers of spindle-shaped cells, Soft matter 10, 2346 (2014).

[5] A. Doostmohammadi and B. Ladoux, Physics of liquid crystals in cell biology, Trends in Cell Biology (2021).

[6] T. B. Saw, A. Doostmohammadi, V. Nier, L. Kocgozlu, S. Thampi, Y. Toyama, P. Marcq, C. T. Lim, J. M. Yeomans, and B. Ladoux, Topological defects in epithelia govern cell death and extrusion, Nature 544, 212 (2017).

[7] O. J. Meacock, A. Doostmohammadi, K. R. Foster, J. M. Yeomans, and W. M. Durham, Bacteria solve the problem of crowding by moving slowly, Nature Physics 17, 205 (2021).

[8] Y. Maroudas-Sacks, L. Garion, L. Shani-Zerbib, A. Livshits, E. Braun, and K. Keren, Topological defects in the nematic order of actin fibres as organization centres of hydra morphogenesis, Nature Physics 17, 251 (2021).

[9] P. Guillamat, C. Blanch-Mercader, G. Pernollet, K. Kruse, and A. Roux, Integer topological defects organize stresses driving tissue morphogenesis, Nature materials 21, 588 (2022).

[10] K. Kawaguchi, R. Kageyama, and M. Sano, Topological defects control collective dynamics in neural progenitor cell cultures, Nature 545, 327 (2017).

[11] K. Copenhagen, R. Alert, N. S. Wingreen, and J. W. Shaevitz, Topological defects promote layer formation in Myxococcus xanthus colonies, Nature Physics 17, 211 (2021).

[12] K. D. Endresen, M. Kim, M. Pittman, Y. Chen, and F. Serra, Topological defects of integer charge in cell monolayers, Soft Matter (2021).

[13] D. J. Cleaver, C. M. Care, M. P. Allen, and M. P. Neal, Extension and generalization of the Gay-Berne potential, Phys. Rev. E 54, 559 (1996).

[14] A. Calderón-Alcaraz, J. Munguía-Valadez, S. I. Hernández, A. Ramírez-Hernández, E. J. Sambriski, and J. A. Moreno-Razo, A bidimensional Gay-Berne calamitic fluid: Structure and phase behavior in bulk and strongly confined systems, Frontiers in Physics 8, 10.3389/fphy.2020.622872 (2021).

[15] J. Stelzer, L. Longa, and H.-R. Trebin, Molecular dynamics simulations of a Gay–Berne nematic liquid crystal: elastic properties from direct correlation functions, The Journal of Chemical Physics 103, 3098 (1995).

[16] C. M. Care and D. J. Cleaver, Computer simulation of liquid crystals, Reports on Progress in Physics 68, 2665 (2005).

[17] W.-J. Rappel and L. Edelstein-Keshet, Mechanisms of cell polarization, Current Opinion in Systems Biology 3, 43 (2017).

[18] B. A. Camley and W.-J. Rappel, Velocity alignment leads to high persistence in confined cells, Physical Review E 89, 062705 (2014).

[19] B. Szabo, G. Szöllösi, B. Gönci, Z. Jurányi, D. Selmeczi, and T. Vicsek, Phase transition in the collective migration of tissue cells: experiment and model, Physical Review E 74, 061908 (2006).

[20] P. Zadeh and B. A. Camley, Picking winners in cell-cell collisions: wetting, speed, and contact, bioRxiv (2022).

[21] A. Szabó, R. Ünnep, E. Méhes, W. Twal, W. Argraves, Y. Cao, and A. Czirók, Collective cell motion in endothelial monolayers, Physical biology 7, 046007 (2010).

[22] M. Versaevel, T. Grevesse, and S. Gabriele, Spatial coordination between cell and nuclear shape within micropatterned endothelial cells, Nature Communications 3, 1 (2012).

[23] R. G. Thakar, Q. Cheng, S. Patel, J. Chu, M. Nasir, D. Liepmann, K. Komvopoulos, and S. Li, Cell-shape regulation of smooth muscle cell proliferation, Biophysical Journal 96, 3423 (2009).

[24] We note that, in principle, increasing ridge height could increase density changes even when driven by motility. This would happen if we were moving from no ridges to a weak ridge height.However, we are not likely to be in a situation where small ridges are a perturbation to no ridges. We see that increasing ridge height does not increase alignment (Fig. 3b), and increasing ridge height decreases crossings (Fig. 6). We thus expect increasing ridge height would suppress density changes if collective motility were driving the density increase at the +1 defect.

[25] G. Duclos, C. Erlenkämper, J.-F. Joanny, and P. Silberzan, Topological defects in confined populations of spindle-shaped cells, Nature Physics 13, 58 (2017).

[26] B. Alric, C. Formosa-Dague, E. Dague, L. J. Holt, and M. Delarue, Macromolecular crowding limits growth under pressure, Nature Physics 18, 411 (2022).

[27] I. F. Rizzuti, P. Mascheroni, S. Arcucci, Z. Ben-Mériem, A. Prunet, C. Barentin, C. Rivière, H. Delanoë-Ayari, H. Hatzikirou, J. Guillermet-Guibert, and M. Delarue, Mechanical control of cell proliferation increases resistance to chemotherapeutic agents, Phys. Rev. Lett. 125, 128103 (2020).

[28] B. I. Shraiman, Mechanical feedback as a possible regulator of tissue growth, Proceedings of the National Academy of Sciences 102, 3318 (2005).

[29] Y. Pan, I. Heemskerk, C. Ibar, B. I. Shraiman, and K. D. Irvine, Differential growth triggers mechanical feedback that elevates hippo signaling, Proceedings of the National Academy of Sciences 113, E6974 (2016).

[30] F. Montel, M. Delarue, J. Elgeti, L. Malaquin, M. Basan, T. Risler, B. Cabane, D. Vignjevic, J. Prost, G. Cappello, et al., Stress clamp experiments on multicellular tumor spheroids, Physical Review Letters 107, 188102 (2011).

[31] X. Yang, D. Bi, M. Czajkowski, M. Merkel, M. L. Manning, and M. C. Marchetti, Correlating cell shape and cellular stress in motile confluent tissues, Proceedings of the National Academy of Sciences 114, 12663 (2017).

[32] C. Leclech, D. Gonzalez-Rodriguez, A. Villedieu, T. Lok, A.-M. Déplanche, and A. I. Barakat, Topography-induced large-scale antiparallel collective migration in vascular endothelium, Nature Communications 13, 1 (2022).

[33] C. Londono, M. J. Loureiro, B. Slater, P. B. Lücker, J. Soleas, S. Sathananthan, J. S. Aitchison, A. J. Kabla, and A. P. McGuigan, Nonautonomous contact guidance signaling during collective cell migration, Proceedings of the National Academy of Sciences 111, 1807 (2014).

[34] H. N. Kim, Y. Hong, M. S. Kim, S. M. Kim, and K.-Y. Suh, Effect of orientation and density of nanotopography in dermal wound healing, Biomaterials 33, 8782 (2012).

[35] K.-H. Nam, P. Kim, D. K. Wood, S. Kwon, P. P. Provenzano, and D.-H. Kim, Multiscale cues drive collective cell migration, Scientific Reports 6, 1 (2016).

[36] J. Petzold and E. Gentleman, Intrinsic mechanical cues and their impact on stem cells and embryogenesis, Frontiers in Cell and Developmental Biology 9, 10.3389/fcell.2021.761871 (2021).

[37] E. Makhija, Y. Zheng, J. Wang, H. R. Leong, R. B. Othman, E. X. Ng, E. H. Lee, L. Tucker-Kellogg, Y. H. Lee, H. Yu, et al., Topological defects govern mesenchymal condensations, offering a morphology-based tool to predict cartilage differentiation, bioRxiv (2022).

[38] J. Folkman and A. Moscona, Role of cell shape in growth control, Nature 273, 345 (1978).

[39] C. S. Chen, M. Mrksich, S. Huang, G. M. Whitesides, and D. E. Ingber, Geometric control of cell life and death, Science 276, 1425 (1997).

[40] N. Gjorevski, M. Nikolaev, T. Brown, O. Mitrofanova, N. Brandenberg, F. DelRio, F. Yavitt, P. Liberali, K. Anseth, and M. Lutolf, Tissue geometry drives deterministic organoid patterning, Science 375, eaaw9021 (2022).

[41] W. Zhu, X. Qu, J. Zhu, X. Ma, S. Patel, J. Liu, P. Wang, C. S. E. Lai, M. Gou, Y. Xu, K. Zhang, and S. Chen, Direct 3d bioprinting of prevascularized tissue constructs with complex microarchitecture, Biomaterials 124, 106 (2017).

[42] K. H. Palmquist, S. F. Tiemann, F. L. Ezzeddine, S. Yang, C. R. Pfeifer, A. Erzberger, A. R. Rodrigues, and A. E. Shyer, Reciprocal cell-ECM dynamics generate supracellular fluidity underlying spontaneous follicle patterning, Cell (2022).

[43] B. A. Camley, Patterning by contraction, Cell (2022).

[44] C. Curantz, R. Bailleul, M. Castro-Scherianz, M. Hidalgo, M. Durande, F. Graner, and M. Manceau, Cell shape anisotropy contributes to self-organized feather pattern fidelity in birds, PLoS Biology 20, e3001807 (2022).

[45] A. F. Marée, V. A. Grieneisen, and P. Hogeweg, The Cellular Potts Model and biophysical properties of cells, tissues and morphogenesis, in Single-cell-based models in biology and medicine (Springer, 2007) pp. 107–136.

[46] J. Singh, A. Pagulayan, B. A. Camley, and A. S. Nain, Rules of contact inhibition of locomotion for cells on suspended nanofibers, Proceedings of the National Academy of Sciences 118, e2011815118 (2021).

[47] J. Gay and B. Berne, Modification of the overlap potential to mimic a linear site–site potential, The Journal of Chemical Physics 74, 3316 (1981).

[48] R. Ananthakrishnan and A. Ehrlicher, The forces behind cell movement, International Journal of Biological Sciences 3, 303 (2007).

[49] G. Duclos, S. Garcia, H. G. Yevick, and P. Silberzan, Perfect nematic order in confined monolayers of spindle-shaped cells, Soft Matter 10, 2346 (2014).

[50] W.-J. Rappel, A. Nicol, A. Sarkissian, H. Levine, and W. F. Loomis, Self-organized vortex state in two-dimensional Dictyostelium dynamics, Physical Review Letters 83, 1247 (1999).

[51] S. Jain, V. M. Cachoux, G. H. Narayana, S. de Beco, J. D’alessandro, V. Cellerin, T. Chen, M. L. Heuzé, P. Marcq, R.-M. Mège, et al., The role of single-cell mechanical behaviour and polarity in driving collective cell migration, Nature Physics 16, 802 (2020).

[52] B. A. Camley and W.-J. Rappel, Physical models of collective cell motility: from cell to tissue, Journal of Physics D: Applied Physics 50, 113002 (2017).

[53] R. Alert and X. Trepat, Physical models of collective cell migration, Annual Review of Condensed Matter Physics 11, 77 (2020).

[54] M. C. Marchetti, J.-F. Joanny, S. Ramaswamy, T. B. Liverpool, J. Prost, M. Rao, and R. A. Simha, Hydrodynamics of soft active matter, Reviews of modern physics 85, 1143 (2013).

[55] M. Nonomura, Study on multicellular systems using a phase field model, PloS one 7, e33501 (2012).

[56] B. A. Camley, Y. Zhang, Y. Zhao, B. Li, E. Ben-Jacob, H. Levine, and W.-J. Rappel, Polarity mechanisms such as contact inhibition of locomotion regulate persistent rotational motion of mammalian cells on micropatterns, Proceedings of the National Academy of Sciences 111, 14770 (2014).

[57] D. A. Kulawiak, B. A. Camley, and W.-J. Rappel, Modeling contact inhibition of locomotion of colliding cells migrating on micropatterned substrates, PLoS Computational Biology 12, e1005239 (2016).

[58] J. Löber, F. Ziebert, and I. S. Aranson, Collisions of deformable cells lead to collective migration, Scientific reports 5, 1 (2015).

[59] Y. Cao, R. Karmakar, E. Ghabache, E. Gutierrez, Y. Zhao, A. Groisman, H. Levine, B. A. Camley, and W.-J. Rappel, Cell motility dependence on adhesive wetting, Soft Matter 15, 2043 (2019).

[60] B. Loewe, M. Chiang, D. Marenduzzo, and M. C. Marchetti, Solid-liquid transition of deformable and overlapping active particles, Physical Review Letters 125, 038003 (2020).

[61] A. Hopkins, M. Chiang, B. Loewe, D. Marenduzzo, and M. C. Marchetti, Local yield and compliance in active cell monolayers, Physical Review Letters 129, 148101 (2022).

[62] G. Zhang, R. Mueller, A. Doostmohammadi, and J. M. Yeomans, Active inter-cellular forces in collective cell motility, Journal of the Royal Society Interface 17, 20200312 (2020).

[63] A. Jayaram, A. Fischer, and T. Speck, From scalar to polar active matter: Connecting simulations with mean-field theory, Physical Review E 101, 022602 (2020).

[64] R. Großmann, I. S. Aranson, and F. Peruani, A particle-field approach bridges phase separation and collective motion in active matter, Nature Communications 11, 1 (2020).

[65] A. M. Menzel and T. Ohta, Soft deformable self-propelled particles, EPL (Europhysics Letters) 99, 58001 (2012).

[66] E. Palsson and H. G. Othmer, A model for individual and collective cell movement in Dictyostelium discoideum, Proceedings of the National Academy of Sciences 97, 10448 (2000).

[67] X. Li, R. Balagam, T.-F. He, P. P. Lee, O. A. Igoshin, and H. Levine, On the mechanism of long-range orientational order of fibroblasts, Proceedings of the National Academy of Sciences 114, 8974 (2017).

[68] B. A. Camley, Y. Zhao, B. Li, H. Levine, and W.-J. Rappel, Crawling and turning in a minimal reaction-diffusion cell motility model: coupling cell shape and biochemistry, Physical Review E 95, 012401 (2017).

[69] I. Nwogbaga and B. A. Camley, Coupling cell shape and velocity leads to oscillation and circling in keratocyte galvanotaxis, Biophysical Journal (2022).

[70] A. R. Singh, T. Leadbetter, and B. A. Camley, Sensing the shape of a cell with reaction diffusion and energy minimization, Proceedings of the National Academy of Sciences 119, e2121302119 (2022).

[71] G. A. Reddy and P. Katira, Differences in cell death and division rules can alter tissue rigidity and fluidization, Soft Matter 18, 3713 (2022).

[72] M. Czajkowski, D. M. Sussman, M. C. Marchetti, and M. L. Manning, Glassy dynamics in models of confluent tissue with mitosis and apoptosis, Soft Matter 15, 9133 (2019).

[73] S. Preibisch, S. Saalfeld, and P. Tomancak, Globally optimal stitching of tiled 3d microscopic image acquisitions, Bioinformatics 25, 1463 (2009).

[74] D. Ershov, M.-S. Phan, J. W. Pylvänäinen, S. U. Rigaud, L. Le Blanc, A. Charles-Orszag, J. R. W. Conway, R. F. Laine, N. H. Roy, D. Bonazzi, G. Duménil, G. Jacquemet, and J.-Y. Tinevez, Trackmate 7: integrating state-of-the-art segmentation algorithms into tracking pipelines, Nature Methods 19, 829 (2022).

